# Physicochemical, functional, and evolutionary characteristics of protein loop regions in human and *Escherichia coli* proteomes

**DOI:** 10.1101/2023.06.09.544332

**Authors:** Lin Zhang, Hafumi Nishi

**Affiliations:** Department of Applied Information Sciences, Graduate School of Information Sciences, Tohoku University, Sendai, Miyagi 980-8579, Japan; Tohoku Medical Megabank Organization, Tohoku University, Sendai, Miyagi 980-8573, Japan; Faculty of Core Research, Ochanomizu University, Tokyo, 112-8610, Japan

**Keywords:** protein loop region, eukaryotic, prokaryotic, secondary structure

## Abstract

Protein loops often play crucial roles in the formation of binding and enzyme active sites. However, the general structural and biological characteristics of these loops remain unclear. In this study, we investigated protein loop regions on a large scale from structural and evolutionary perspectives. After removing redundancy at the protein chain level, 555,516, 102,901, and 24,818 loops were extracted from the entire PDB, Homo sapiens, and *Escherichia coli* proteins, respectively. Regardless of whether they were isolated from humans or *E. coli*, numerous loop sequences tended to be unique among proteins or protein chains. However, loop properties exhibited high similarity or conservation, including length, distance, and stretch. The CATH classification analysis suggested that most loops connected the same superfamily, while the heterogeneity of superfamily context repertoires was not explained at the topology or homologous level. In contrast, the functions of conserved loops between human and *E. coli* proteins were not consistently conserved, with sequences exhibiting considerable divergence in enrichment. The amino acid composition profiles showed that loops from humans exhibited a preference for serine, whereas *E. coli* loops had biases for glycine and alanine. Although the amino acid composition was primarily determined by the species, the composition of certain special types of loops clustered separately from other classes, suggesting the existence of conserved loops with complex functions. Collectively, this study provides a detailed overview of protein loops from the structural, functional, and evolutionary perspectives and a vast natural loop repertoire for mining additional information.

## Introduction

Protein loops play critical roles in protein function. In particular, the loops on protein surfaces often participate in enzyme reactions and ligand/protein binding as they are exposed to solvents. Although the loop regions of homologous proteins may display higher variability in terms of sequence composition, they also exhibit conserved architectures to some extent ^1^. Exchanging loop sequences for proteins can enable the gain of function ^2–4^. Notably, the reconstruction of humanized antibodies or the creation of antibody-mimetic proteins supports loop sequence functions ^5–7^.

Loops also contribute to the evolution of protein functions. Loop regions generate variability through diverse sequence compositions, leading to new or altered functions ^8^. Sequence changes in loop regions are common during the evolution of enzyme functions ^9^. In fact, considering the structural and sequence (dis)similarity of loop regions is a more efficient means of classifying proteins or characterizing protein evolution compared to conventional approaches (e.g., measuring conserved protein core regions) ^10^. Accordingly, many studies have sought to elucidate the roles of the protein loop region in protein function and evolution. Typically, these studies have focused on loop structural and physicochemical properties: loop sequence, number of residues in a loop, local loop structure, and distance and stretch (normalized distance) between the endpoints of a loop ^2, 6, 11–16^. For instance, in contrast to the loop stretch, Choi et al. demonstrated that the loop distance distribution appears to be independent of the number of residues, while highly stretched loops are easier to predict than contracted loops ^17^.

The amino acid composition of protein loops has also been investigated from the functional characteristic perspective. For example, phenylalanine is overabundant (e.g., glycine) in hotspot loops, exhibiting a high average binding energy; thus, hot loops are commonly used to recognize protein targets ^18^. Omega loops, often associated with regulatory functions and biomolecular recognition ^19–21^, are biased toward specific amino acids, such as glycine, proline, tyrosine, aspartate, serine, and asparagine ^19, 22^. Moreover, eukaryotic proteins have evolved myriad notable features compared to prokaryotic proteins, including longer sequences ^23–26^ and abundant multi-domain proteins ^27–29^. Amino acid frequencies have been studied in-depth within intrinsically disordered regions, where the difference in disordered regions between eukaryotic and prokaryotic proteins is caused by a shift in the frequencies of serine, proline, and isoleucine ^30^. However, to the best of our knowledge, the analysis of loop properties on a larger scale and their comparative analysis between eukaryotes and prokaryotes from an evolutionary perspective has not been conducted. Moreover, the amino acid frequencies of large-scale loops from eukaryotic and prokaryotic proteins are less understood.

In this study, we aim to describe and compare the properties of protein loops acquired from the entire protein data bank (PDB), Homo sapiens, and *Escherichia coli* (*E. coli*) and assess the loop structures and functions from an evolutionary perspective. By characterizing the three-loop properties, i.e., length, distance, and stretch, we demonstrate the level of similarity among humans and *E. coli* datasets. In particular, similar loop distances and stretch distributions were observed, as well as longer loops in humans compared to *E. coli*. Loop functions identified based on CATH and Pfam annotations show that protein loops are highly capable of gaining or losing unique functions in humans or *E. coli*. The preferences for specific amino acids vary between eukaryotes and prokaryotes. For instance, there were differences in serine, glycine, and alanine frequencies between humans and *E. coli*.

## Materials and Methods

### Loop definition

A protein loop structure was defined as any region classified as “B,” “S,” “T,” or space in the secondary structure assignment. The following additional criteria were applied to curate the data: (1) the number of residues of a loop ranged from 4–50; (2) terminal residues were not located in disordered regions; (3) the loop stretches were no more than 1.2; (4) loop sequences without unknown residues (shown as “X” in the PDB FASTA file); (5) the protein chain termini were skipped when extracting loop regions. For NMR structures, the first model was used for further analysis.

### Dataset construction

Three-dimensional structures and secondary structure assignments with missing protein region information were downloaded from protein data bank (PDB, December 2019). A non-redundant chain dataset (the redundancy among chains was removed with > 90% sequence identity (S90)) served as the primary dataset, however, redundant (the entire PDB) and S100 (removing redundant sequences based on the 100% identity threshold) datasets were also assessed (Figure 1). Protein chains were further divided into different datasets according to the source organisms: the entire PDB, Homo sapiens, and *E. coli*. Note that protein chains with multiple source organisms were removed. For the non-redundant S90 datasets, 55,654, 12,212, and 2,334 protein chains from the entire PDB, Homo sapiens, and *E. coli*, respectively, were used separately for loop extraction. The number of protein chains was 417,925, 103,984, and 21,958 for the redundant datasets and 85,029, 20,283, and 4,143 for the non-redundant S100 datasets.

**Figure 1.**
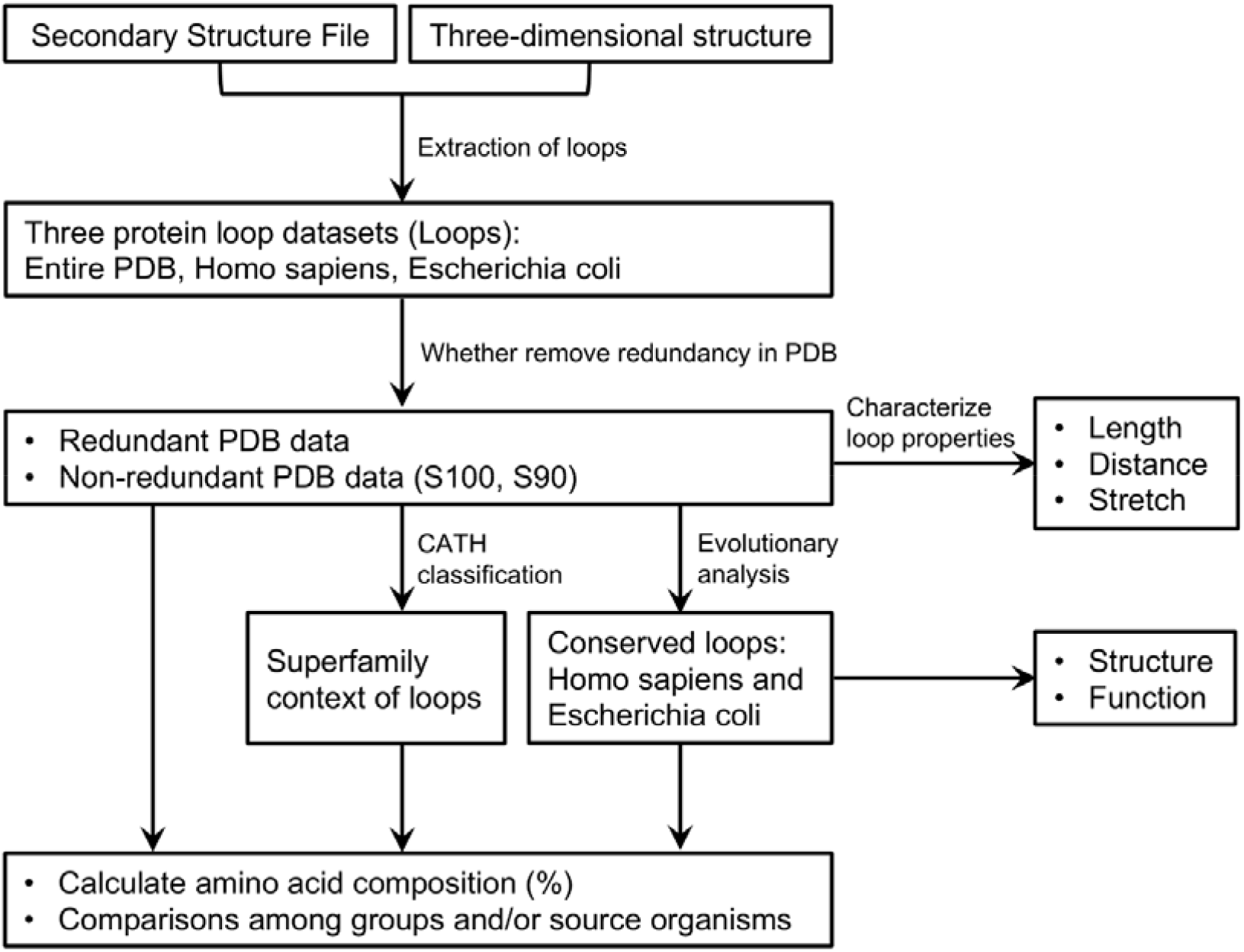
Workflow of this study.

### Loop properties

Three loop properties were used: loop length, loop distance, and loop stretch. Loop length is the number of residues in the loop. The loop distance (*l*) is defined as the distance between the first and last C alpha atoms of the loop. The loop stretch (l) is the normalized loop distance, l = *l/l_max_*. The maximum distance *l*_max_ is associated with the number of residues (n) and calculated using the formulas g(n/2 − 1) + d and g(n − 1)/2 for even and odd n, respectively, where g = 6.046 and d = 3.46 Å ^17, 31, 32^. The loop stretch indicates how stretched a loop is. Ideally, the loop stretch for a fully stretched loop should be 1. However, because the values of g and d are theoretical approximations, it has been reported that the loop stretch values can be > 1 ^31^.

### Structural and functional domain annotations for loops

The structural domains of the loops were assigned using the CATH classifications (Figure 1). Specifically, combined with the SIFTS database (structure integration with function, taxonomy, and sequence), the CATH v4.2 and CATH daily update file (latest classifications, 2020/01/13) were used to annotate superfamilies of both terminal residues of a loop.

From an evolutionary perspective, conserved loop sequences between humans and *E. coli* were identified. They were then used to analyze the structures and functions using CATH classifications and the Pfam database. The CATH classification was similar to that used in the above analysis. Regarding Pfam analysis, the corresponding loops of humans and *E. coli* were obtained according to the conserved loop sequences. The Pfam IDs of the terminal residues of a loop were extracted from the SIFTS files. Eventually, Pfam clans were extracted based on Pfam IDs.

### Superfamily context repertoire

The CATH protein structure classification database was applied to provide information on the evolutionary relationships of the protein domains; those sharing evolutionary similarities were assigned to the same CATH superfamily ^33^. Considering the CATH superfamily, loops can link a common or diverse superfamily. Note that only the superfamilies of the terminal residues of a loop were considered. As loop sequences may be duplicated, each redundant loop sequence possesses a superfamily context repertoire that composed of all superfamilies associated with different loops that share the same sequence, regardless of whether they are located in other regions of the same protein chain or in different molecules. Superfamily context repertoires can be homogeneous or heterogeneous, depending on whether the loops share the same superfamily context.

### Amino acid composition profiles

The amino acid compositions of the protein loop sequences from different datasets, i.e., different organisms, sub-datasets of loops connecting common or diverse superfamily contexts, and conserved loops, were calculated. Clustered heat maps were generated using Ward’s variance minimization algorithm, and the distance was computed using Euclidean.

### Identification of orthologous-like sequences for *Archaeoglobus fulgidus* dataset analysis

BLASTp version 2.13.0 was used to identify orthologous-like sequences in the human and *E. coli.* datasets. Selected amino acid sequences of *Archaeoglobus fulgidus* were used as a query sequence and searched against PDB. To obtain the best matched sequence, an e-value threshold of 1e-2 and a minimum query coverage of 50% were applied.

## Results

### Protein loop extraction and statistics

We downloaded protein secondary and 3D structure data from the PDB in December 2019. The numbers of protein chains used in this study are listed in Table 1. Combined with the SIFTS database, we identified protein loops and calculated loop features, namely length, distance, stretch, amino acid sequence, and connected secondary structure. The first model was based on NMR structures. Multiple loops on a single protein molecule were identified and analyzed separately. Subsequently, we assigned structural and functional annotations to loops by detecting the CATH superfamily of the loop terminal residues. The numbers of loops, unique loop sequences, loops annotated by the CATH database, and percentages of non-redundant loop sequences are shown in Table 1. Loops from the entire PDB (28.98%), humans (31.37%), and *E. coli* (24.41%) were not annotated by CATH after removing redundant sequences in the PDB with 90% sequence identity (S90). In this study, a “loop” indicates a single loop region on a protein structure, while a “loop sequence” indicates the amino acid sequence of the loop regions. Therefore, two different loops in different protein structures can have the same loop amino acid sequence. Compared to the total number of loops, the number of unique loop sequences decreased by 24.21%, 21.93%, and 10.77% in the PDB, human, and *E. coli* datasets, respectively. Intriguingly, the non-redundant loop sequences were extremely high for the S90 datasets, up to 91.75% in *E. coli*. and 85.13% in the human dataset. The higher percentage of non-redundant loop sequences in *E. coli* may be due to a preference for specific loops in human protein or biases of data depositions in the PDB.

**Table 1.**
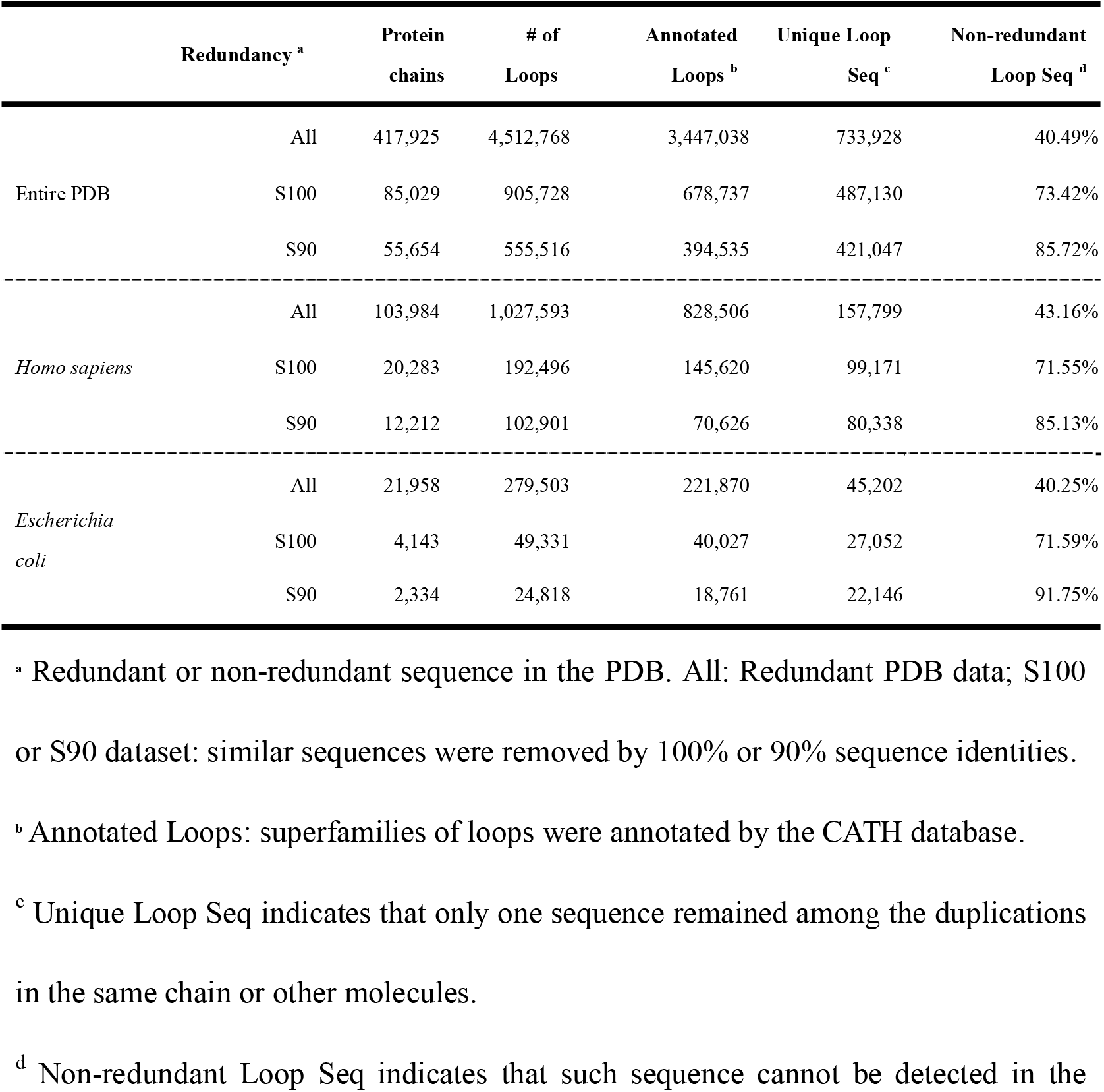
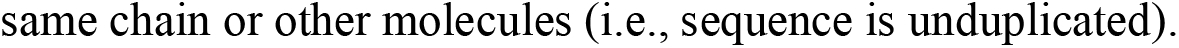
Protein loop statistics.

|  | Redundancy <sup>a</sup> | Protein chains | # of Loops | Annotated Loops <sup>b</sup> | Unique Loop Seq <sup>c</sup> | Non-redundant Loop Seq <sup>d</sup> |
| --- | --- | --- | --- | --- | --- | --- |
|  | All | 417,925 | 4,512,768 | 3,447,038 | 733,928 | 40.49% |
| Entire PDB | S100 | 85,029 | 905,728 | 678,737 | 487,130 | 73.42% |
|  | S90 | 55,654 | 555,516 | 394,535 | 421,047 | 85.72% |
| ----- |  |  |  |  |  |  |
|  | All | 103,984 | 1,027,593 | 828,506 | 157,799 | 43.16% |
| *Homo sapiens* | S100 | 20,283 | 192,496 | 145,620 | 99,171 | 71.55% |
|  | S90 | 12,212 | 102,901 | 70,626 | 80,338 | 85.13% |
| ----- |  |  |  |  |  |  |
|  | All | 21,958 | 279,503 | 221,870 | 45,202 | 40.25% |
| *Escherichia coli* | S100 | 4,143 | 49,331 | 40,027 | 27,052 | 71.59% |
|  | S90 | 2,334 | 24,818 | 18,761 | 22,146 | 91.75% |

<sup>a</sup> Redundant or non-redundant sequence in the PDB. All: Redundant PDB data; S100 or S90 dataset: similar sequences were removed by 100% or 90% sequence identities.
<sup>b</sup> Annotated Loops: superfamilies of loops were annotated by the CATH database.
<sup>c</sup> Unique Loop Seq indicates that only one sequence remained among the duplications in the same chain or other molecules.
<sup>d</sup> Non-redundant Loop Seq indicates that such sequence cannot be detected in the
same chain or other molecules (i.e., sequence is unduplicated).

### Comparisons of protein loop properties among the entire PDB, Homo sapiens, and *Escherichia coli*

We first investigated three main loop structural properties: loop length, distance, and stretch (Figure 2). The loop length frequency distribution ranged from 4–50. Predictably, we found that the frequencies decreased as the loop length and percentage of short loops (four and five residues) increased in *E. coli*. The loops on the protein structure are shown in Figure 2A, and the unique loop sequences are shown in Figure 2B. A larger difference in the frequency of 4 and 5 residue loops was observed between humans and *E. coli*, while the other loop lengths were similar. The loop distances increased as the number of loop residues increased, while the loop stretches exhibited the opposite trend (Figure 2C and 2D). We also observed the loop distance and stretch distributions (Figure 2E and 2F). Two peaks in the loop distance distribution (5–6 Å and 8–10 Å) and one in the loop stretch distribution (∼0.6) were observed.

**Figure 2.**
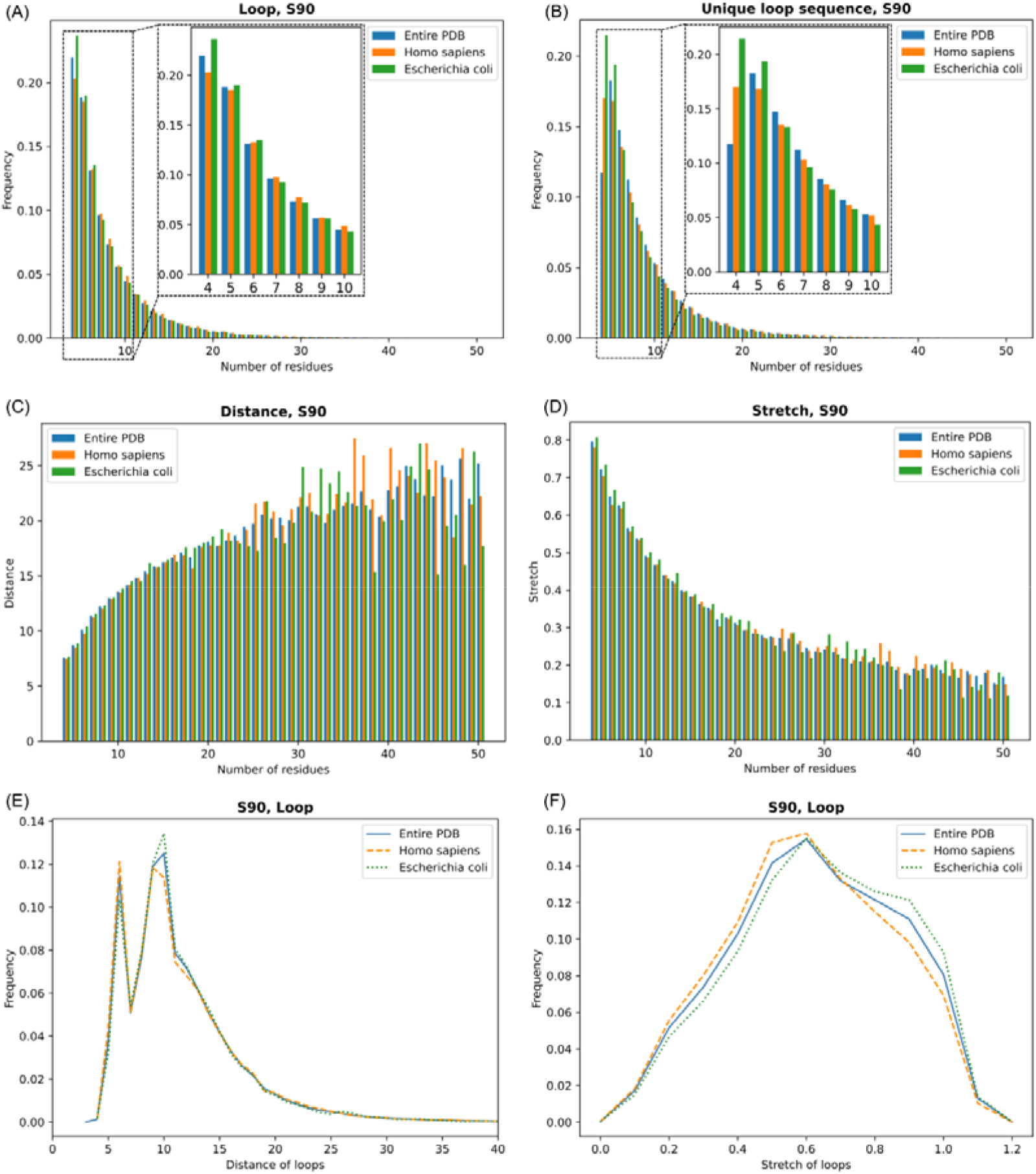
Comparison of protein loop characteristics among the entire PDB, *Homo sapiens*, and *Escherichia coli*. (A)–(B) Frequency distribution of protein loops with the residue number ranging from 4–50. The frequency was calculated according to individual loops and unique loop sequences. The loop distance (C) and stretch (D) of loops containing different residues. The Y axis in (C) and (D) represents average distance and stretch values in terms of length, respectively. Loop distance (E) and stretch (F) frequency distribution.

To investigate whether loop distance and stretch are related to loop length, we analyzed their distributions using loops containing 8, 10, 12, 14, and 16 residues from the entire PDB, human, and *E. coli* datasets (Figure S1A–S1F). The distance distributions were not altered for each dataset when the number of residues increased, whereas the loop stretch distributions shifted back compared to the distances. In addition, quantile-quantile plots (Q–Q plots), a method for evaluating the similarity between two probability distributions by comparing their quantiles, showed that the human and *E. coli* loop distance distributions were from the same probability distribution (Figure S1G and S1F). Altogether, these observations suggest that loop distance may be independent of residue number, loop stretching may not. Moreover, they are not unique to certain organisms, indicating that these properties are evolutionarily conserved.

To explain the two peaks in the loop distance distributions, we examined the linked secondary structure types and calculated the amino acid composition of the corresponding loops. A tendency for the linked secondary structure types was observed. That is, compared to the peak at approximately 8–10 Å, that at 5–6 Å exhibited a preference for linking two strands (strand-strand) compared to the other connection types (helix-helix, helix-strand, and strand-helix). In total, 71.90%, 77.80%, and 67.88% loops with distances of 5–6 Å from the entire PDB, human, and *E. coli* datasets were found to link strand-strands, respectively. Therefore, these shorter distance loops are used to connect the antiparallel β strands. Regarding the amino acid composition of the loops from all datasets, the 5–6 Å loops preferred glycine, while the 8–10 Å peak contained more leucine residues. The relationships between amino acid composition and loop stretch were also investigated; the loops from the entire PDB, human, and *E. coli* datasets were divided into 12 stretch bins with a 0.1 difference for the neighboring bins (Figure S2). For each amino acid, we further calculated the variance of the 12 stretch bins. Findings indicated that amino acids, such as cysteine, methionine, histidine, tryptophan, arginine, and lysine, contributed less to loop stretching than aspartate, glycine, and proline. See Figure S2 for full details.

### Structural and functional analysis based on CATH classification

Next, we analyzed the protein loop structures and functions based on the evolutionary relationships of the protein domains using the CATH database. The four main levels of the CATH classification hierarchy are class (C), architecture (A), topology/fold (T), and the homologous superfamily (H). In addition, we detected the superfamilies of the terminal residues of a loop as the “context” of the loop. Therefore, the superfamily context of a loop was described as a combination of two superfamily IDs. The S90 datasets of the entire PDB, human, and *E. coli* were used here. If the two superfamily IDs assigned to the loop’s two ends are A and B, then all extracted loops can be divided into four groups: A=B, A!=B, A, or B, and null (Figure 3A). A=B indicates that loops connect an identical superfamily, whereas A!=B represents loops connected to different superfamilies. In groups A and B, only one of the terminal residues are annotated with a superfamily, whereas the null group contains no terminal residues found in a superfamily. Most loops were successfully annotated by CATH as a superfamily for both terminal residues. To confirm whether loops linking common or diverse superfamilies were related to loop length, we assessed the loop length distributions of the A=B and A!=B groups using loops from the entire PDB, human, and *E. coli* S90 datasets (Figure 3B–D). In comparing loops among different organisms, the loops linking identical superfamilies showed similar distribution patterns of loop length, however, the distributions of loops linking diverse superfamilies were not, and they were deferred to the A=B group.

**Figure 3.**
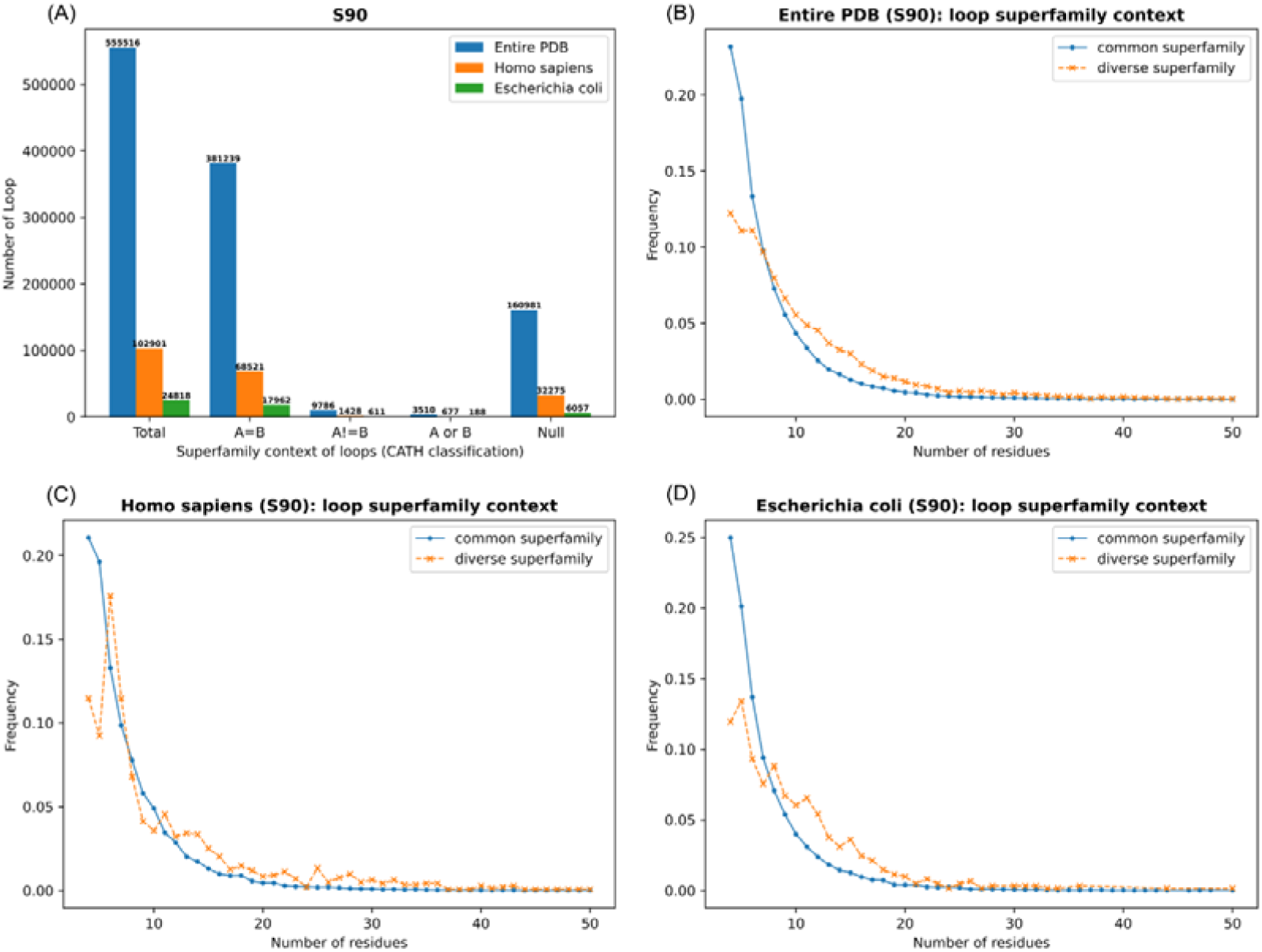
Superfamily context of loops using the CATH database. (A) CATH classification of loops. A and B represent the superfamilies of terminal residues. Four groups were generated based on a loop connecting common, diverse, or no superfamilies. (B)–(D) The loop length distributions for loops connecting common and diverse superfamilies in the entire PDB, Homo sapiens, and *E. coli* datasets.

Although the context analyses were performed for individual loops, different loops may share the same loop amino acid sequence. Therefore, to investigate the effect of amino acid sequences on the superfamily context of loops, we investigated and compared the superfamily context of each loop sequence from different loops. Since the same loop sequence may come from different proteins, the superfamily context repertoire of a loop sequence can be homogeneous (a loop sequence from different loops shares a common superfamily context) or heterogeneous (multiple superfamily contexts were observed). The loops used in this analysis were filtered using the following criteria: (1) the loop should share its sequence with other loops, and (2) both terminal residues of the loop should have the CATH annotation. A total of 41,668, 7,688, and 1, 251 loop sequences were obtained from the entire PDB, Homo sapiens, and *E. coli*, respectively. The filtered loop sequences were then assigned to either homogeneous or heterogeneous repertoires by gradually reducing the CATH levels from 4 to 1 (from top to bottom; four levels are C, A, T, and H). The number of loop sequences divided into homogeneous or heterogeneous repertoires is shown in Table 2. Many heterogeneous repertoires were found, indicating that such loop sequences can be involved in multiple superfamily contexts. When we increased the CATH levels, the number of heterogeneous repertoires was predicted to decrease as fewer structural or functional constraints were considered. However, the number of heterogeneous repertoires did not decrease considerably with higher CATH levels, suggesting that the heterogeneity was not explained at the topology or homologous superfamily level of CATH classifications. We also examined the loop length distributions of the homogeneous and heterogeneous repertoire loop sequences (Figure S3). Consequently, the number of loops with four residues was extremely high for heterogeneous repertoires. This might be due to the short length of loops containing four residues making them highly likely to be observed in multiple domains or superfamilies.

**Table 2.** Superfamily context repertoire of loops from S90 non-redundant datasets.

|  | # of unique loop sequences | # of redundant loop sequences | CATH_levels <sup>a</sup> | Homogeneous | Heterogeneous |
| --- | --- | --- | --- | --- | --- |
| Entire PDBs | 301,462 | 41,668 | (C, A, T, H) | 22,102 | 19,566 |
|  |  |  | (C, A, T) | 22,860 | 18,808 |
|  |  |  | (C, A) | 24,069 | 17,599 |
|  |  |  | (C) | 29,496 | 12,172 |
| *Homo sapiens* | 55,277 | 7,688 | (C, A, T, H) | 6,455 | 1,233 |
|  |  |  | (C, A, T) | 6,514 | 1,174 |
|  |  |  | (C, A) | 6,608 | 1,080 |
|  |  |  | (C) | 6,915 | 773 |
| *Escherichia coli* | 16,948 | 1,251 | (C, A, T, H) | 922 | 329 |
|  |  |  | (C, A, T) | 961 | 290 |
|  |  |  | (C, A) | 1,001 | 250 |
|  |  |  | (C) | 1,125 | 126 |

<sup>a</sup>This analysis was performed considering different CATH levels. Consider the superfamily ID 1.10.510.10 as an example. 1.10.510.10: (C, A, T, H); 1.10.510: (C, A, T); 1.10: (C, A); 1: (C).

### Understanding conserved loops between humans and *E. coli*

In the previous analyses, we separately analyzed human and *E. coli* datasets and compared the results. Here, we detected the conserved loop sequences between humans and *E. coli* according to 100% sequence identity to investigate the structures and functions of protein loops from an evolutionary perspective. In total, 3,350, 2,037, and 1,633 conserved loop sequences were identified in the redundant (all) and non-redundant datasets (S100 and S90), respectively. The conserved loop sequences from the S90 datasets were further analyzed. Compared to the total loop sequences of human and *E. coli* datasets, 2.03% of human and 7.37% of *E. coli* loop sequences were conserved between the two species. In terms of individual loops, 3.27% of human and 10.54% of *E. coli* loops were related to conserved loop sequences. The results suggested that the conserved loop sequences were overrepresented in protein chains from both humans and *E. coli* but even more overrepresented in *E. coli*, as the percentage differences between unique loop sequences and individual loops of *E. coli* (3.17%) was much larger than that of humans (1.24%) (i.e., compared to humans, the conserved sequence tended to be duplicated in *E. coli*). The top 30 conserved loop sequences from humans and *E. coli* are listed in Table S1 according to the ranking of the corresponding loop numbers of each sequence. The abundance of each conserved loop sequence differed between humans and *E. coli*.

After characterizing the conserved loops, we examined their functional annotations using CATH and Pfam databases. The length distributions of individual loops associated with the conserved loop sequences are shown in Figure S4A. Regarding the CATH classifications, similar results were found in that many loops failed to be annotated by CATH, and most successfully annotated loops were found to connect an identical superfamily (Figure S4B). However, the conserved loop may be the result of (1) evolutionary conservation following the divergence of a protein loop from a common ancestral sequence; (2) convergence on the same loop sequence in unrelated proteins; (3) recombination or convergence if the same sequence is present at different locations in the same superfamily ^1, 34–37^. Considering that we found that most conserved loops were assigned to the A=B group, this may indicate common ancestry of the proteins as they come from the same superfamily, while the loops could be either (1) or (3) of the above, suggesting a mix of homologous and convergent cases. Note that only a few conserved loops were detected in the A!=B group (Figure S4B, 22 from humans and 26 from *E. coli*), which might help to understand the case of obvious convergence. The CATH classification analysis on the A!=B group suggested that the conserved loop sequence tended to show a similar superfamily context in unrelated proteins within or between species (Table S2). Table S2 provides insights regarding whether the same structural/functional solution has been discovered by evolution from independent starting points or if these are merely coincidences.

In addition, the top ten conserved loops of the CATH superfamily and Pfam clan are displayed in Figure S5, and their functional descriptions are listed in Table S3. Comparing humans and *E. coli*, several fundamental functions were conserved. For the CATH database, the conserved functions were immunoglobulins, P-loop-containing nucleotide triphosphate hydrolases, NAD(P)-binding Rossmann-like domain, periplasmic binding protein-like II, and glutaredoxin. For the Pfam database, the conserved functions were P-loop_NTPase, periplasmic binding proteins (PBPs), NADP_Rossmann, beta _propeller, and thioredoxin. However, many functions were not conserved. These observations indicate that although the loop sequences were conserved, it was highly possible to gain or lose unique functions in humans and *E. coli*.

To better understand and compare the functions of conserved loop sequences, we acquired ligands/chemical components of related PDB entries and analyzed their ligand-binding sites in humans and *E. coli*. We found that not all conserved loops participated in ligand binding. Moreover, the functions of the conserved loop sequence can also be conserved; that is, for a given conserved loop sequence, the loop regions from human and *E. coli* PDB entries were observed to potentially bind not only to the same ligand but also to completely different ligands or only one of the loop regions was identified as a ligand. To visualize these three conditions, the conserved loop sequences, VAEH, ADNSG, and GPNMGG, were chosen as examples (Figure 4). Each loop sequence of the human proteins (top column) was identical to that of the corresponding *E. coli* protein (bottom). For VAEH, the phosphate ion (PO_4_) interacted with residue E44 of 2Y1H (chain A) of humans, and pyridoxal-5′-phosphate (PLP) was found to bind to residue H119 of 4LW2 (chain A) of *E. coli*. For ADNSG, guanosine-5′-diphosphate (GDP) bound to residues A48 and S51 of 5G53 (chain C) in humans, but no ligand was observed in 6S0K (chain L) of *E. coli*. For GPNMGG, the same ligand, adenosine-5′-diphosphate (ADP), was identified, which binds to M1137 and G1139 of 2O8B (chain B) in humans compared to M617 and G619 of 6I5F (chain A) in *E. coli*. Specifically, we assume that this loop (GPNMGG) is a P-loop, as it showed the same feature as that reported previously ^38^; that is the structural P-loop contains a four-residue backbone fragment (i.e., the GXXX pattern) that primarily connects a strand and helix. Additionally, these loops (2O8B and 6I5F) were classified into the 3.40.50.300 CATH superfamily, which is described as a P-loop containing nucleotide triphosphate hydrolases.

**Figure 4.**
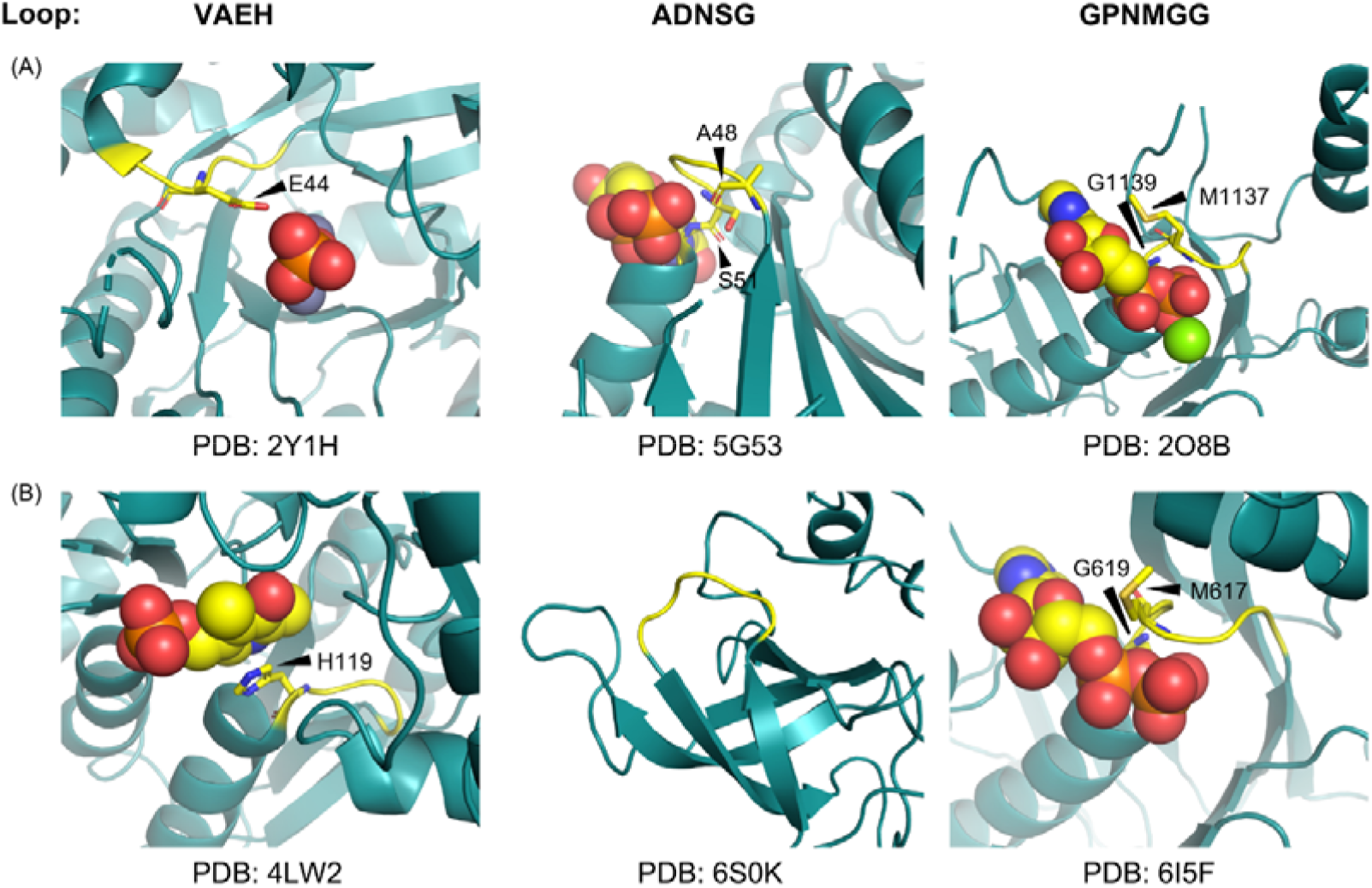
Examples of conserved loop sequences between humans and *E. coli*. Visualization of the ligand-binding sites of (A) humans and (B) *E. coli* were generated using PyMol. Three conserved loop sequences are shown. The top (human) sequence was the same as that from the corresponding bottom *(E. coli)*. The protein loop regions are marked in yellow, and the remaining regions are represented in dark green. The binding sites are shown as sticks, and each residue is labeled with one letter. Ligands are shown as balls.

### Amino acid composition and comparison among various groups with different resource organisms

To examine the contributions of amino acids to individual loops and their properties, we calculated and compared amino acid composition profiles among different species, superfamily combinations, and repertoire groups. The loops in the PDB, human, and *E. coli* datasets were divided into six categories: all loops, all unique loop sequences, A=B, A!=B, homogeneous, and heterogeneous groups. In addition, the conserved unique loop sequences and their related loops within 3D structures of proteins from humans and *E. coli* were analyzed. As shown in Figure 5, a clustered heatmap was used to visualize the similarities and differences in amino acid composition profiles in different groups (the amino acid composition ratios for each group are shown in Table S4). As a result, various loop categories from different organisms were separated, suggesting that the amino acid differences among organisms were larger than those within an organism. Interestingly, A!=B, heterogeneous and conserved loops exhibited opposite trends, as each category among organisms was closely related. Specifically, because the loop number of the A!=B category was low (Figure 3A), we assumed that only a few loops could connect different CATH superfamily members, and these loops may tend to be conserved between humans and *E. coli*. Likewise, the number of loops with a heterogeneous superfamily context repertoire was small (Table 2), and they may also be conserved. Collectively, we hypothesized that few loops that showed more “complex” functions (e.g., connecting various superfamilies or involved in heterogeneous superfamily context) might tend to be conserved.

**Figure 5.**
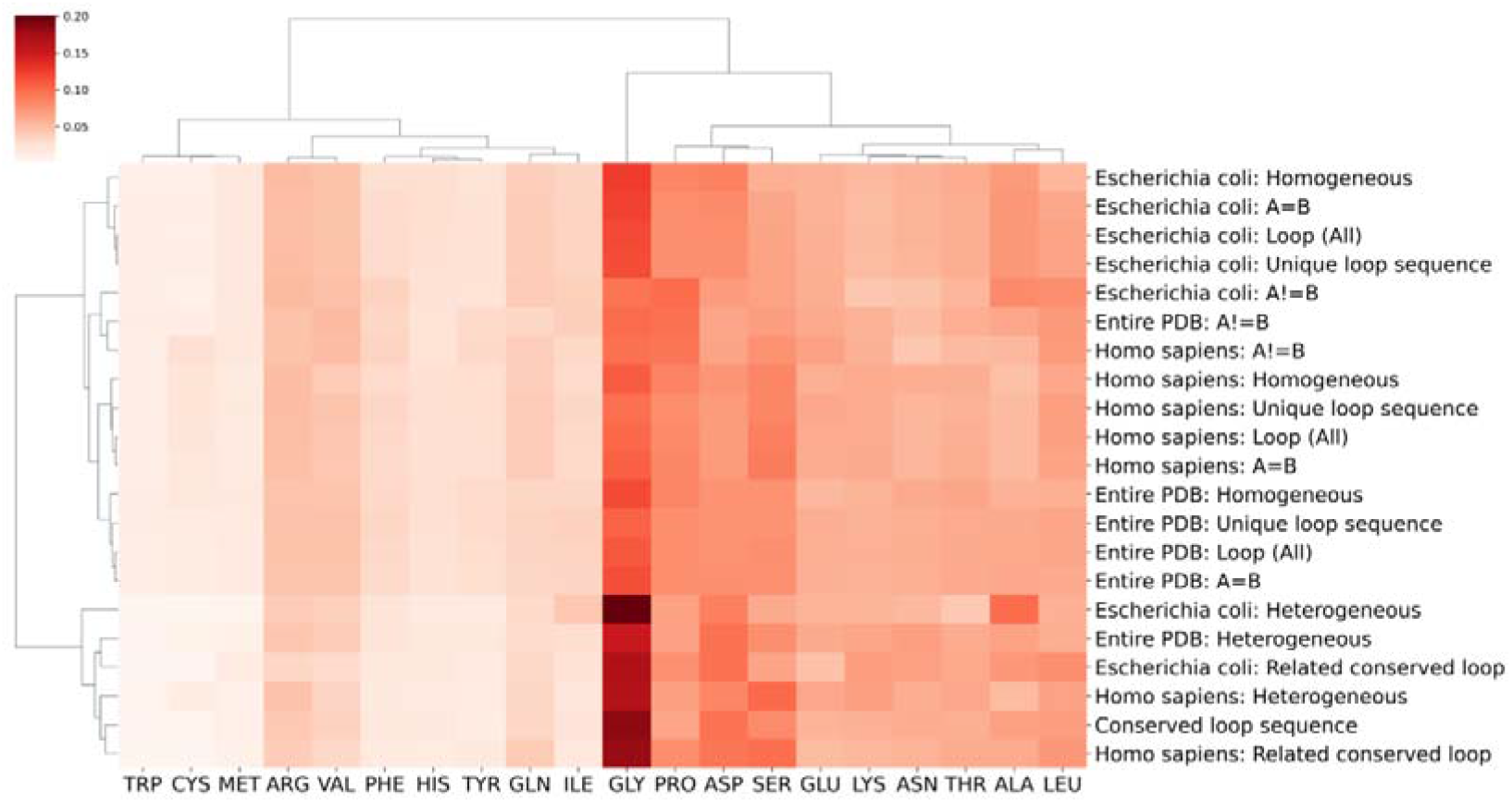
Clustered heatmap of amino acid composition profiles of loops from various groups with different resource organisms. Protein loops from three resource organisms are included: Entire_PDBs, humans, and *E. coli*. The Loop (all) and Unique loop sequence categories represent loop sequences acquired based on individual loops and unique loop sequences, respectively. The A=B and A!=B categories represent loops linking common and diverse superfamilies. Homogeneous and heterogeneous categories were the heterogeneity of the superfamily context of loops.

To elucidate the factors affecting the unique amino acid composition distributions of the different groups, we compared each amino acid composition profile among groups (among categories and resource organisms, Figure 6). Each amino acid composition that differed by > 1.5% was considered to be the main contributor to the differences ^30^. In terms of categories (Figure 6A and B), the amino acid composition distributions of all loops from humans and *E. coli* were highly similar to those of all and unique sequences. Compared to similar distributions between A=B and A!=B in humans, the A=B category in *E. coli* prefers glycine, while the A!=B category prefers leucine and proline. For both humans and *E. coli*, the homogeneous category showed a preference for proline over glycine in the heterogeneous category. Notably, the conserved loops from both humans and *E. coli* were biased toward glycine and aspartic acid. In terms of resource organisms (Figure 6C), we observed that loops (all loops and unique loop sequences) in humans displayed preferences for serine compared to that of glycine and alanine in *E. coli*. This may be due to serine serving as a target of serine/threonine kinases prevalent in eukaryotes ^39^. One potential reason for the abundance of glycine and alanine in *E. coli* is that the cost of producing alanine and glycine (also serine) is the lowest ^40^. We further examined the frequencies of glycine, alanine, and serine in humans and *E. coli* (Figure S6) and found that glycine and alanine were decreased in all datasets in humans compared to *E. coli* and that the serine frequency exhibited the opposite trend. Moreover, the frequency differences in glycine, alanine, and serine were widespread and not unique to particular datasets.

**Figure 6.**
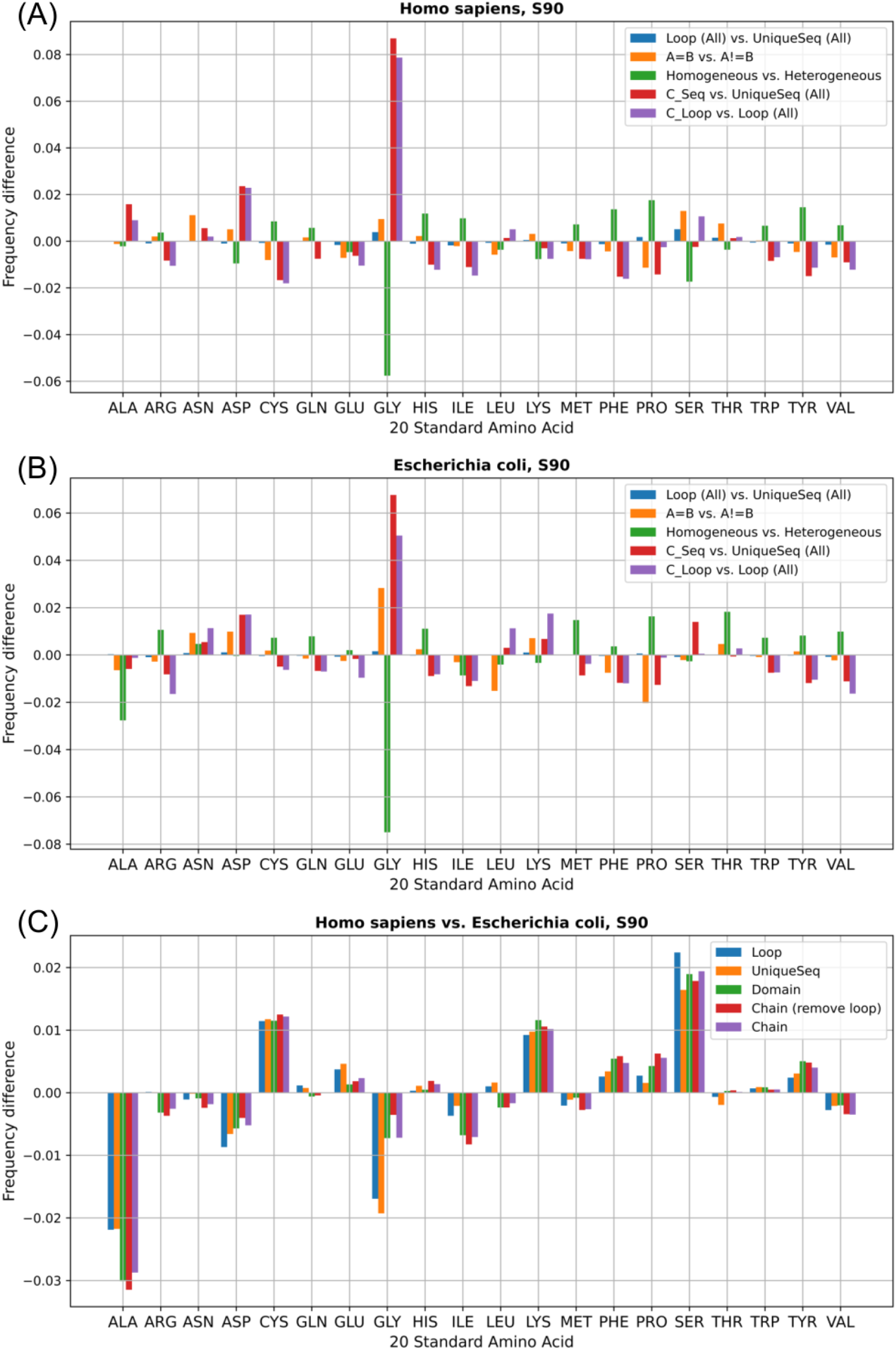
Amino acid frequency differences among groups and source organisms. Comparisons among groups from (A) Homo sapiens and (B) *Escherichia coli.* Positive values represent amino acids showing a higher frequency in the front categories and negative values represent a higher frequency in the latter categories. E.g., for loop (all) vs. UniqueSeq (all), frequency difference = loop (all) - UniqueSeq (all). (C) Comparison of amino acid composition between humans and *E. coli*. The *E. coli* frequencies are used as a baseline; that is, positive values indicate that the amino acids are more abundant in humans. Domains: Domain regions of protein chains that form a loop. Chains: entire protein chains with a loop. Compared to Chains, Chain_remove_loop indicates that the loop regions were removed, and only the remaining segments of protein chains were used.

## Discussion

Loop properties of various membranes and soluble proteins have been studied previously. In particular, Choi et al. ^17^ reported that the loop distance distribution patterns appear to be independent of the loop length, whereas the loop stretch is not. We observed similar results in a large-scale loop dataset comprising different organisms, suggesting that these findings were not unique to a specific species and that such loops tend to be conserved during evolution. Additionally, compared to Choi’s studies, we found two similar peaks (5–6 Å and 8–10 Å) in the loop distributions and determined that the 5–6 Å distances strongly preferred antiparallel beta loops. In this study, we further analyzed their amino acid composition to elucidate the underlying mechanisms of such preferences. Loops with distances of 5–6 Å from the PDB, human, and *E. coli* datasets exhibited a preference for glycine. One possible reason for this is that glycine is the smallest amino acid and might contribute to fitness of short distances. In particular, we observed that loops with distances of 8–10 Å did not show any preference for the linked secondary structure types but had a bias toward leucine. However, the reasons for these preferences remain unknown. Furthermore, the distances of long loops containing 31–50 residues were considerably larger in the human dataset than in the *E. coli* dataset (Mann–Whitney U-test, *p* = 0.0429). Their corresponding stretches were also larger in humans, but not significantly (Mann–Whitney U-test, *p* = 0.1366). These observations indicate that long loops from *E. coli* (> 30 residues) might tend to contract.

In general, the loop stretch is assumed to be < 1. However, as certain parameters used for calculating the loop stretch are empirical values, the loop stretches can be > 1. Therefore, we also examined loops where the stretches were > 1.2. Only a few loops with a stretch of more than 1.2 were detected, specifically 20 loops from humans and one from *E. coli* (Table S5). Therefore, discarding the loops with stretches > 1.2 would not affect the performance of our analyses. Meanwhile, given the small number of loops, it was difficult to estimate the contributions of amino acids to the loops with a larger stretch (Table S6). Therefore, we hypothesized that the higher stretches were due to the improper 3D structures deposited in the PDB.

Loop sequence compositions exhibit high variability in homologous proteins with conserved architectures ^1^. In fact, loop sequences often change during enzyme evolution ^9^. In this study, we found that more than 85% of the loop sequences from the entire PDB, human, and *E. coli* were observed only once, indicating that most of the loops were unique at the protein chain level. Moreover, several loop sequences were detected multiple times and were involved in homogeneous or heterogeneous superfamily context repertoires. These loop sequences may contribute to the variability in the sequence compositions of proteins and chains.

Generally, our observations suggest that certain loop features are robust during the evolutionary process. However, eukaryotes and prokaryotes have their own loop biases. For example, the abundance of conserved loop sequences between humans and *E. coli* differ. The top sequence for humans is QPGGS, which was detected in 102 loops compared to NKHMNADTD in humans, which was found in 38 loops (Table S1). In addition, the VAEH and ADNSG loops formed different three-dimensional conformations and bound (or do not bind) different ligands in human and *E. coli* proteins (Figure 4). Combined with previous studies of the conformational ensemble of loops and the associated dynamics ^41, 42^, our examples provide important insights regarding the mechanisms of conformational loop selection, which might inform the prediction of accurate positions for various loop fragments ^43^.

Intriguingly, we found that protein loops exhibited preferences for specific amino acids, such as glycine, proline, aspartic acid, and leucine. The abundances were widespread among various loop sets (i.e., different categories in this study) and organisms, excluding serine and alanine, which were particularly enriched in humans and *E. coli*, respectively (Table S4). Although human and *E. coli* protein loops similarly favored certain amino acids, their preferences for loop sequences were extremely divergent (Table S1). In addition, many loops from categories or organisms have different biases toward particular amino acids. We assumed that this might be partially associated with post-translational modifications of loops. For example, the potential reason for serine enrichment in humans is that serine is a phosphorylation target, whereas between 1/3 and 2/3 of the eukaryotic proteome are phosphorylated ^44, 45^. Conserved loops showed a preference for glycine and aspartic acid (both are relatively small), suggesting that these two amino acids may be essential for the flexibility and solvent exposure of loops.

In addition, to gain a comprehensive understanding of properties between eukaryotic and prokaryotic proteomes, we characterized the loop properties in *Saccharomyces cerevisiae* and *Archaeoglobus fulgidus* as additional species for comparison. Compared to entire PDBs, human, and *E. coli* datasets, similar findings were observed (Figure S7 and S8). The amino acid compositions of yeast loops exhibit higher similarity with human sequences compared to *E. coli* (Figure S9), as expected. Additionally, archaea displayed unique features, including a preference for short loops and a particularly long distance of loops containing 25 or 28 residues (Figure S8). We speculated that these observations might be due to species-specific features or the small dataset size (only 151 protein chains in archaea; Figure S8). To explore this, we ran BLAST with the 151 sequences from archaea to identify orthologous-like sequences in human and *E. coli* and characterize the loop properties. We found 56 and 42 matched chains in human and *E. coli*, respectively, and the corresponding loops displayed similarity and dissimilarity to human and *E.coli*, including a similar preference for short loops and different amino acid compositions (Figure S10). Unfortunately, due to the small dataset size, the cause of dissimilarity observed in archaea is difficult to explain.

Furthermore, we examined missing regions in a loop to investigate the effect of intrinsically disordered regions (IDRs). We calculated the percentages of loops that include at least one disorder site or fully disordered loops and found that the relative proportions were small among the species, suggesting that the IDRs might not have a considerable effect on our results (Table S7).

In conclusion, this study provides an in-depth understanding of loop structural properties, functions, and evolution on a large scale. It also provides a vast natural loop repertoire for mining additional information, such as the mechanisms for controlling functions and protein folding. We performed analyses using all PDB entries, human, and *E. coli.* proteins, which is considered to be sufficient to describe the details of the loop properties and functions, as well as basic structural analyses. However, since this study is dependent on the database, certain biases may have occurred due to the data deposition. For example, if the depth and width of protein analysis differ considerably between humans and *E. coli*, limitations might arise when assessing the differences between humans and *E. coli*. To improve the performance, we removed redundant protein sequences; however, certain biases may be difficult to remove. Therefore, a more complete analysis of loop structures and their conformational changes is needed. These analyses could contribute to the discovery of further characteristics of naturally emerging and designed proteins and finer predictions of robust protein structures.

## Acknowledgments

The authors thank Prof. Kengo Kinoshita and Prof. Matsuyuki Shirota for valuable discussions. This study was financially supported by the Development of Key Technologies for the Next-Generation Artificial Intelligence/Robots of the New Energy and Industrial Technology Development Organization (NEDO).

## Author Contributions

H.N. designed the research. L.Z. performed the experiments. L.Z. and H.N. analyzed the data. L.Z. and H.N. prepared the manuscript.

## Supplementary Information

**Figure S1. Distributions of distance and stretch for loops with different lengths.**

Loop distance (A–C) and stretch (D–F) distributions for loops containing certain numbers of residues, 8, 10, 12, 14, and 16 residues. (G–H) Quantile-quantile (Q–Q) plot, for evaluating the similarity between two probability distributions by comparing their quantiles, was used to compare the distance and stretch distributions of humans and *E. coli*.

**Figure S2. Amino acid compositions for each stretch bin.**

Loops were obtained from (A) entire PDB, (B) Homo sapiens, and (C) *Escherichia coli*. (A–C) The upper panel represents amino acid distribution and the lower panel represents the corresponding variance. (D) Number of loops in 12 stretch bins.

**Figure S3. Length distribution of loops with homogeneous and heterogeneous superfamily context repertoires.**

The analysis was performed using loops from the (A) entire PDB, (B) Homo sapiens, and (C) *Escherichia coli* datasets.

**Figure S4. General understanding of loop length and CATH classifications of conserved loops.**

(A) Conserved loop length distribution in humans and *Escherichia coli*. (B) CATH classification of the conserved loops from humans and *E. coli*.

**Figure S5. Top ten CATH superfamily and Pfam clan.**

(A) and (B) represented the top ten CATH superfamilies and Pfam clans of conserved loops between humans and *E.coli*, respectively.

**Figure S6. Frequency of glycine, alanine, and serine in five datasets from humans and *E. coli*.**

The datasets included only specific segments of amino acid sequences and were Domain (sequences of protein domain regions), Chain (sequences of whole protein chains), Chain_RMLoop (sequences of protein chains with loop region removed), UniqueSeq (all unique loop sequences), and Loop (all loop sequences, which indicated loop sequences are redundant).

**Figure S7. Characterizing properties of *Saccharomyces cerevisiae* protein loops.**

(A) Protein loop statistics. (B) Visualizations of loop properties. (C) Superfamily context of a loop based on the CATH database. (D) Clustered heatmap of amino acid composition profiles. (E) Amino acid frequency differences among groups. (F) Statistics of superfamily context repertoire of loops from S90 non-redundant datasets.

**Figure S8. Characterizing properties of *Archaeoglobus fulgidus* protein loops.**

(A) Protein loop statistics. (B) Visualizations of loop properties. (C) Superfamily context of a loop based on the CATH database. (D) Clustered heatmap of amino acid composition profiles. (E) Amino acid frequency differences among groups. (F) Statistics of superfamily context repertoire of loops from S90 non-redundant datasets.

**Figure S9. Amino acid frequency comparisons between *Saccharomyces cerevisiae*, human, and *E. coli*.**

The human and E. coli frequencies were used as a baseline; that is, positive values indicate that the amino acids are more abundant in yeast.

**Figure S10. Blast analysis using 151 protein chains from *Archaeoglobus fulgidus* against human and *E. coli* proteins.**

(A) and (B) represent the loop property characterizations using matched protein chains from human and *E. coli*, respectively. The following parameters were used to select the best matched amino acid sequence in the blast analysis: e value 1e-2 and the coverage of the query sequence (*Archaeoglobus fulgidus* chains) is ≥ 50%.

**Table S1. Top 30 conserved loop sequences from humans and *E. coli*.**

**Table S2. CATH classifications of the conserved loops in the A!=B group (S90).**

**Table S3. Descriptions of the top ten CATH superfamily and Pfam clan between humans and *E.coli*.**

**Table S4. Twenty standard amino acid compositions (ratios) of loops from various groups (S90).**

**Table S5. Loops with stretch values > 1.2.**

**Table S6. Amino acid compositions of loops with larger stretches (> 1.2)**

**Table S7. Disorder statistics among the datasets.**

## Notes

### Competing Interest Statement

The authors have declared no competing interest.

## References

1. Papaleo E, Saladino G, Lambrughi M, Lindorff-Larsen K, Gervasio FL, Nussinov R. The Role of Protein Loops and Linkers in Conformational Dynamics and Allostery. Chem Rev. 2016;116(11):6391–6423. doi:10.1021/acs.chemrev.5b00623

2. Pardon E, Haezebrouck P, De Baetselier A, et al. A Ca2+-binding Chimera of Human Lysozyme and Bovine α-Lactalbumin That Can Form a Molten Globule *. J Biol Chem. 1995;270(18):10514–10524. doi:https://doi.org/10.1074/jbc.270.18.10514

3. Wolfson AJ, Kanaoka M, Lau FTK, Ringe D. Insertion of an elastase-binding loop into interleukin-1β. Protein Eng Des Sel. 1991;4(3):313–317. doi:10.1093/protein/4.3.313

4. Toma S, Campagnoli S, Margarit I, et al. Grafting of a calcium-binding loop of thermolysin to Bacillus subtilis neutral protease. Biochemistry. 1991;30(1):97–106. doi:10.1021/bi00215a015

5. Ito T, Nishi H, Kameda T, et al. Combination Informatic and Experimental Approach for Selecting Scaffold Proteins for Development as Antibody Mimetics. Chem Lett. 2021;50(11):1867–1871.

6. Queen C, Schneider WP, Selick HE, et al. A humanized antibody that binds to the interleukin 2 receptor. Proc Natl Acad Sci. 1989;86(24):10029 LP - 10033. doi:10.1073/pnas.86.24.10029

7. Riechmann L, Clark M, Waldmann H, Winter G. Reshaping human antibodies for therapy. Nature. 1988;332(6162):323-327. doi:10.1038/332323a0

8. Furnham N, Sillitoe I, Holliday GL, et al. Exploring the Evolution of Novel Enzyme Functions within Structurally Defined Protein Superfamilies. PLOS Comput Biol. 2012;8(3):e1002403. https://doi.org/10.1371/journal.pcbi.1002403

9. Nestl BM, Hauer B. Engineering of flexible loops in enzymes. Acs Catal. 2014;4(9):3201–3211. doi:doi.org/10.1021/cs500325p

10. Panchenko AR, Madej T. Structural similarity of loops in protein families: toward the understanding of protein evolution. BMC Evol Biol. 2005;5(1):10. doi:10.1186/1471-2148-5-10

11. Fernandez-Fuentes N, Oliva B, Fiser A. A supersecondary structure library and search algorithm for modeling loops in protein structures. Nucleic Acids Res. 2006;34(7):2085–2097. doi:10.1093/nar/gkl156

12. Hildebrand PW, Goede A, Bauer RA, et al. SuperLooper—a prediction server for the modeling of loops in globular and membrane proteins. Nucleic Acids Res. 2009;37(suppl_2):W571–W574. doi:10.1093/nar/gkp338

13. Wojcik J, Mornon JP, Chomilier J. New efficient statistical sequence-dependent structure prediction of short to medium-sized protein loops based on an exhaustive loop classification 11Edited by J. M. Thornton. J Mol Biol. 1999;289(5):1469–1490. doi:https://doi.org/10.1006/jmbi.1999.2826

14. Leszczynski JF, Rose GD. Loops in globular proteins: a novel category of secondary structure. Science (80-). 1986;234(4778):849 LP - 855. doi:10.1126/science.3775366

15. Ring CS, Kneller DG, Langridge R, Cohen FE. Taxonomy and conformational analysis of loops in proteins. J Mol Biol. 1992;224(3):685–699. doi:https://doi.org/10.1016/0022-2836(92)90553-V

16. Choi Y, Deane CM. FREAD revisited: Accurate loop structure prediction using a database search algorithm. Proteins Struct Funct Bioinforma. 2010;78(6):1431–1440. doi:https://doi.org/10.1002/prot.22658

17. Choi Y, Agarwal S, Deane CM. How long is a piece of loop? PeerJ. 2013;1:e1. doi:10.7717/peerj.1

18. Gavenonis J, Sheneman BA, Siegert TR, Eshelman MR, Kritzer JA. Comprehensive analysis of loops at protein-protein interfaces for macrocycle design. Nat Chem Biol. 2014;10(9):716–722. doi:10.1038/nchembio.1580

19. Mager PP, Walther H. A hydrophilic omega-loop (Tyr181 to Tyr188) in the nonsubstrate binding area of HIV-1 reverse transcriptase. Drug Des Discov. 1996;14(3):225–239.

20. Fetrow JS. Omega loops; nonregular secondary structures significant in protein function and stability. FASEB J. 1995;9(9):708–717.

21. Rose GD. Loops in globular proteins: Identification of a novel category of secondary structure. Published online 1986.

22. Espadaler J, Querol E, Aviles FX, Oliva B. Identification of function-associated loop motifs and application to protein function prediction. Bioinformatics. 2006;22(18):2237–2243.

23. Apic G, Gough J, Teichmann SA. Domain combinations in archaeal, eubacterial and eukaryotic proteomes. J Mol Biol. 2001;310(2):311–325.

24. Gerstein M, Levitt M. Comprehensive assessment of automatic structural alignment against a manual standard, the scop classification of proteins. Protein Sci. 1998;7(2):445–456.

25. Liu J, Rost B. CHOP proteins into structural domain like fragments. Proteins Struct Funct Bioinforma. 2004;55(3):678–688.

26. Ekman D, Björklund ÅK, Frey-Skött J, Elofsson A. Multi-domain proteins in the three kingdoms of life: orphan domains and other unassigned regions. J Mol Biol. 2005;348(1):231–243.

27. Ekman D, Björklund ÅK, Elofsson A. Quantification of the elevated rate of domain rearrangements in metazoa. J Mol Biol. 2007;372(5):1337–1348.

28. Gerstein M. How representative are the known structures of the proteins in a complete genome? A comprehensive structural census. Fold Des. 1998;3(6):497–512.

29. Apic G, Gough J, Teichmann SA. An insight into domain combinations. Bioinformatics. 2001;17(suppl_1):S83–S89.

30. Basile W, Salvatore M, Bassot C, Elofsson A. Why do eukaryotic proteins contain more intrinsically disordered regions? PLOS Comput Biol. 2019;15(7):e1007186. https://doi.org/10.1371/journal.pcbi.1007186

31. Tastan O, Klein-Seetharaman J, Meirovitch H. The Effect of Loops on the Structural Organization of α-Helical Membrane Proteins. Biophys J. 2009;96(6):2299–2312. doi:https://doi.org/10.1016/j.bpj.2008.12.3894

32. Flory PJ, Volkenstein M. Statistical mechanics of chain molecules. Published online 1969.

33. Sillitoe I, Dawson N, Lewis TE, et al. CATH: expanding the horizons of structure-based functional annotations for genome sequences. Nucleic Acids Res. 2019;47(D1):D280–D284. doi:10.1093/nar/gky1097

34. Lupas AN, Ponting CP, Russell RB. On the Evolution of Protein Folds: Are Similar Motifs in Different Protein Folds the Result of Convergence, Insertion, or Relics of an Ancient Peptide World? J Struct Biol. 2001;134(2):191–203. doi:https://doi.org/10.1006/jsbi.2001.4393

35. Makarova KS, Grishin N V. The Zn-peptidase superfamily: functional convergence after evolutionary divergence11Edited by J. M. Thornton. J Mol Biol. 1999;292(1):11–17. doi:https://doi.org/10.1006/jmbi.1999.3059

36. Todd AE, Orengo CA, Thornton JM. Evolution of function in protein superfamilies, from a structural perspective11Edited by A. R. Fersht. J Mol Biol. 2001;307(4):1113–1143. doi:https://doi.org/10.1006/jmbi.2001.4513

37. Blouin C, Butt D, Roger AJ. Rapid evolution in conformational space: A study of loop regions in a ubiquitous GTP binding domain. Protein Sci. 2004;13(3):608–616. doi:https://doi.org/10.1110/ps.03299804

38. Kinoshita K, Sadanami K, Kidera A, Go N. Structural motif of phosphate-binding site common to various protein superfamilies: all-against-all structural comparison of protein–mononucleotide complexes. Protein Eng Des Sel. 1999;12(1):11–14. doi:10.1093/protein/12.1.11

39. Leonard CJ, Aravind L, Koonin E V. Novel families of putative protein kinases in bacteria and archaea: evolution of the “eukaryotic” protein kinase superfamily. Genome Res. 1998;8(10):1038–1047.

40. Akashi H, Gojobori T. Metabolic efficiency and amino acid composition in the proteomes of Escherichia coli and Bacillus subtilis. Proc Natl Acad Sci. 2002;99(6):3695–3700.

41. Boehr DD, Nussinov R, Wright PE. The role of dynamic conformational ensembles in biomolecular recognition. Nat Chem Biol. 2009;5(11):789–796.

42. Gu Y, Li DW, Bru schweiler R. Decoding the mobility and time scales of protein loops. J Chem Theory Comput. 2015;11(3):1308–1314.

43. de Oliveira SHP, Shi J, Deane CM. Building a Better Fragment Library for De Novo Protein Structure Prediction. PLoS One. 2015;10(4):e0123998. https://doi.org/10.1371/journal.pone.0123998

44. Vlastaridis P, Kyriakidou P, Chaliotis A, Van de Peer Y, Oliver SG, Amoutzias GD. Estimating the total number of phosphoproteins and phosphorylation sites in eukaryotic proteomes. Gigascience. 2017;6(2). doi:10.1093/gigascience/giw015

45. Cohen P. The origins of protein phosphorylation. Nat Cell Biol. 2002;4(5):E127–E130. doi:10.1038/ncb0502-e127

