## Supplementary Information for "Physicochemical, functional, and evolutionary characteristics of protein loop regions in human and *Escherichia coli* proteomes"

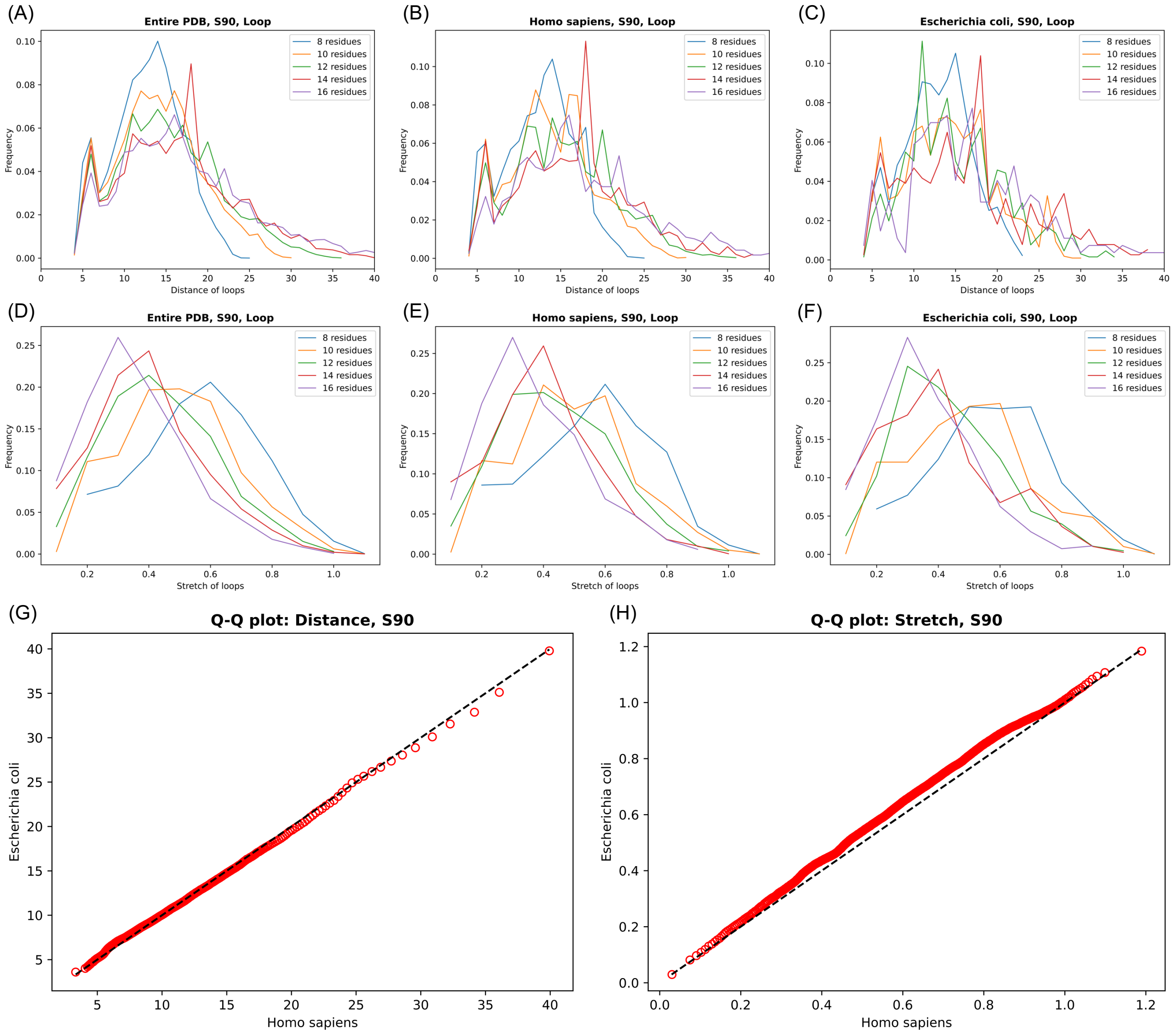


**Figure S1. Distributions of distance and stretch for loops with different loop lengths.**

Loop distance (A–C) and stretch (D–F) distributions for loops containing certain numbers of residues, 8, 10, 12, 14, and 16 residues. (G–H) Quantile-quantile (Q–Q) plot, for evaluating the similarity between two probability distributions by comparing their quantiles, was used to compare the distance and stretch distributions of humans and *E. coli*.

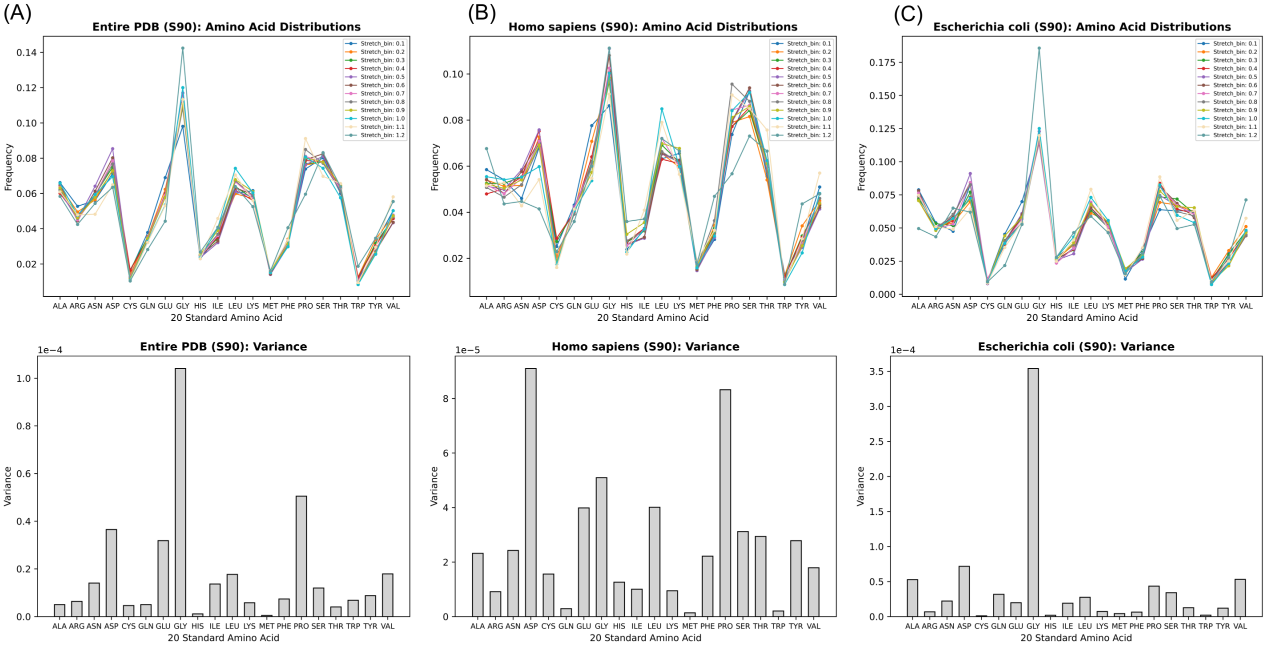


(D)

| | Entire PDB | Homo sapiens | Escherichia coli |
|---|---:|---:|---:|
| Stretch_bin_0.1 | 2,656 | 584 | 121 |
| Stretch_bin_0.2 | 19,145 | 3,812 | 710 |
| Stretch_bin_0.3 | 34,926 | 6,817 | 1,429 |
| Stretch_bin_0.4 | 46,928 | 9,605 | 1,875 |
| Stretch_bin_0.5 | 71,159 | 14,236 | 2,915 |
| Stretch_bin_0.6 | 86,545 | 16,730 | 3,691 |
| Stretch_bin_0.7 | 76,513 | 14,225 | 3,667 |
| Stretch_bin_0.8 | 73,510 | 13,441 | 3,285 |
| Stretch_bin_0.9 | 59,041 | 9,980 | 2,804 |
| Stretch_bin_1.0 | 64,072 | 10,146 | 3,324 |
| Stretch_bin_1.1 | 19,613 | 3,127 | 926 |
| Stretch_bin_1.2 | 1,408 | 198 | 71 |
| Total loops | 555,516 | 102,901 | 24,818 |

**Figure S2. Amino acid compositions for each stretch bin.**
Loops were obtained from (A) entire PDB, (B) Homo sapiens, and (C) *Escherichia coli*. (A–C) The upper panel represents amino acid distribution and the lower panel represents the corresponding variance. (D) Number of loops in 12 stretch bins.

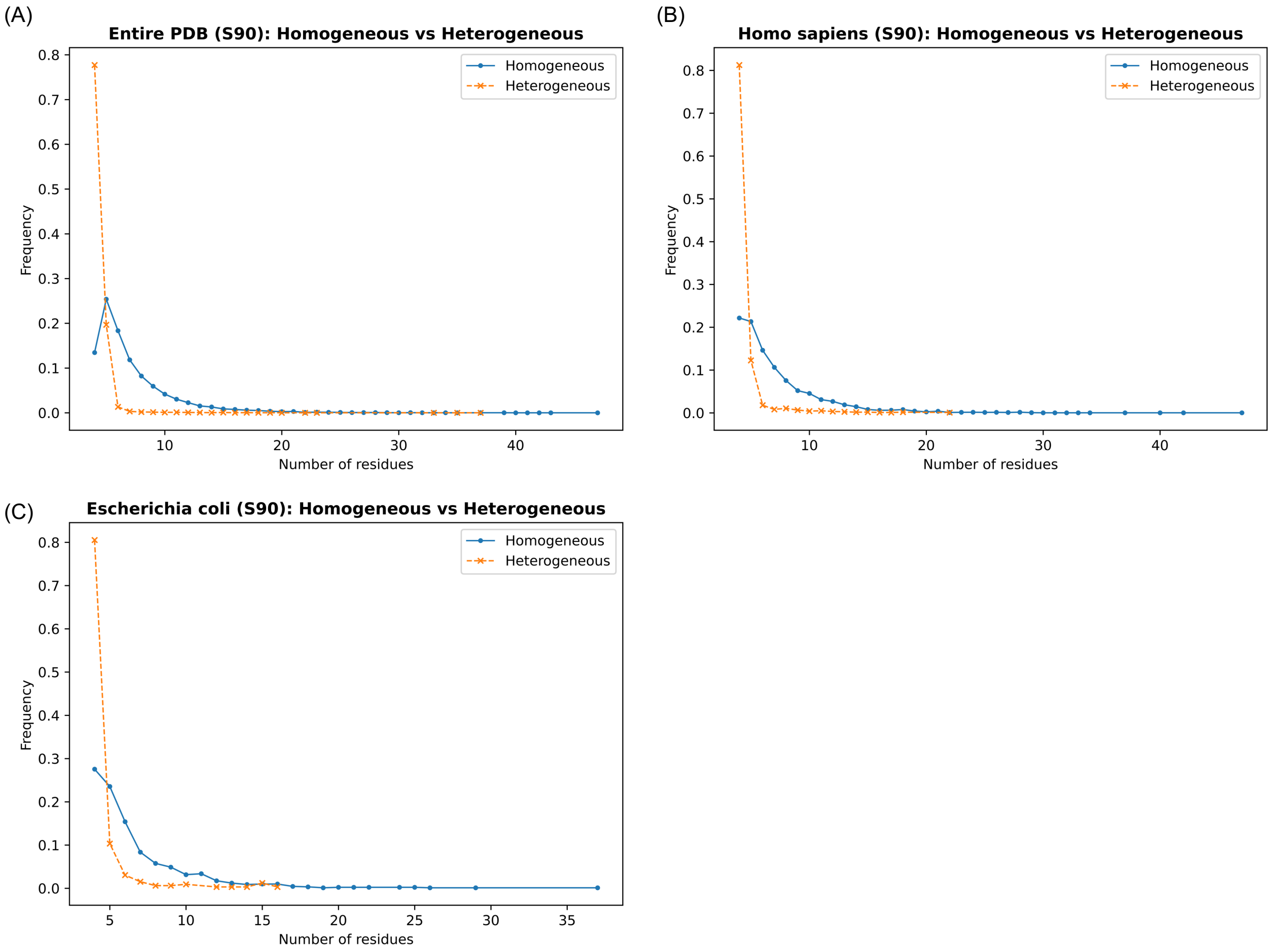


**Figure S3. Length distribution of loops with homogeneous and heterogeneous superfamily context repertoires.**
The analysis was performed using loops from the (A) entire PDB, (B) Homo sapiens, and (C) *Escherichia coli* datasets.

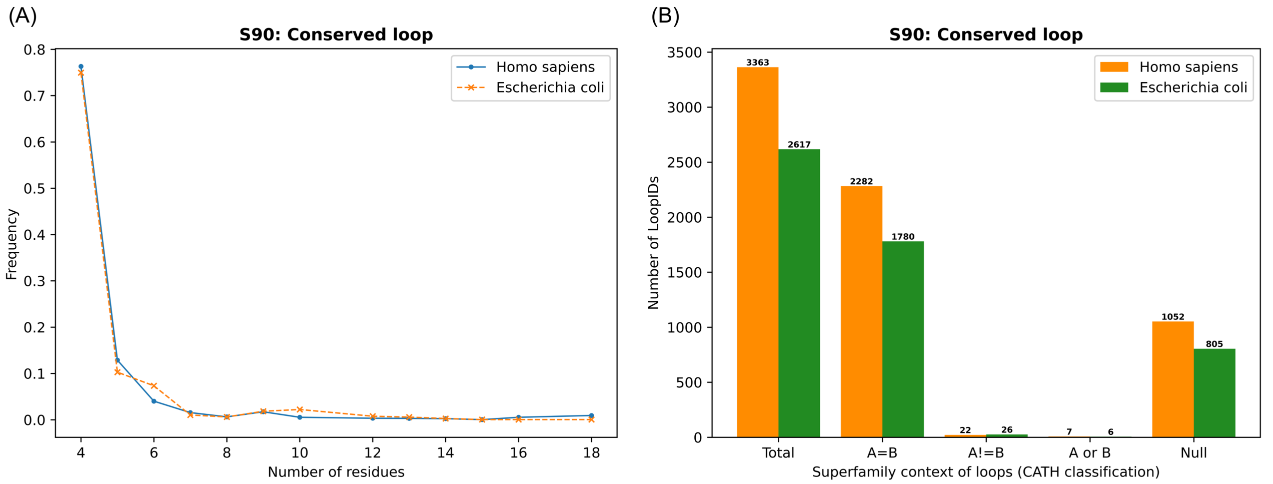


**Figure S4. General understanding of loop length and CATH classifications of conserved loops.**
(A) Conserved loop length distribution in humans and *Escherichia coli*. (B) CATH classification of the conserved loops from humans and *E. coli*.

(A)

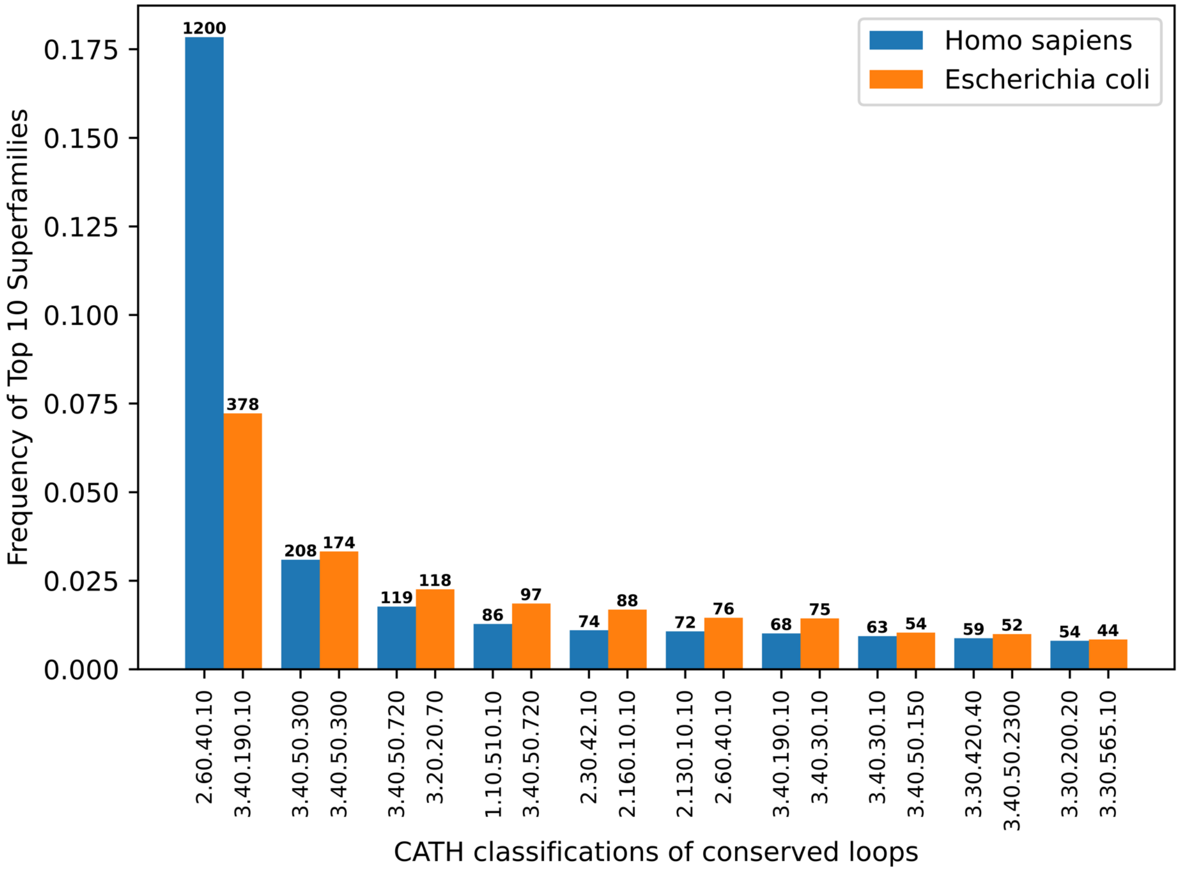


(B)

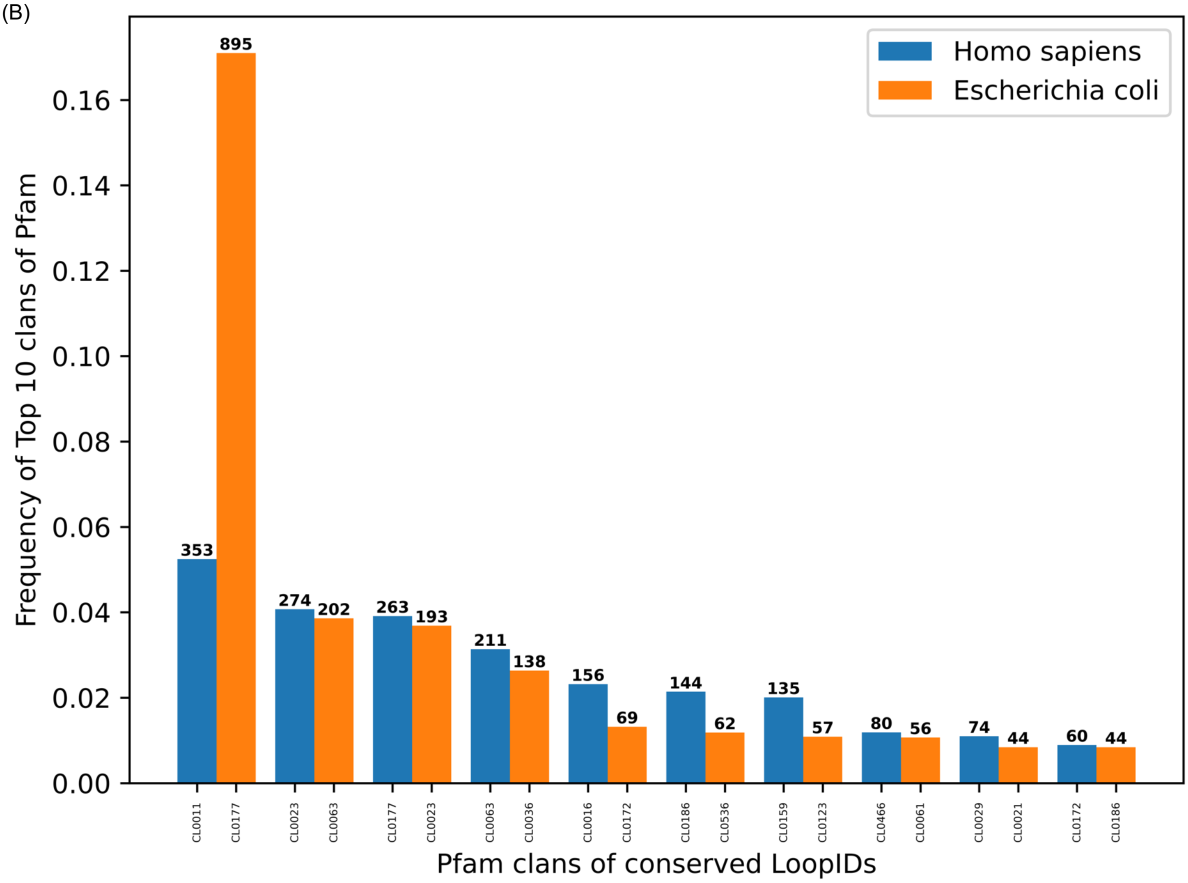


**Figure S5. Top ten CATH superfamily.**
(A) and (B) represented the top ten CATH superfamilies and Pfam clans of conserved loops between humans and E.coli, respectively.

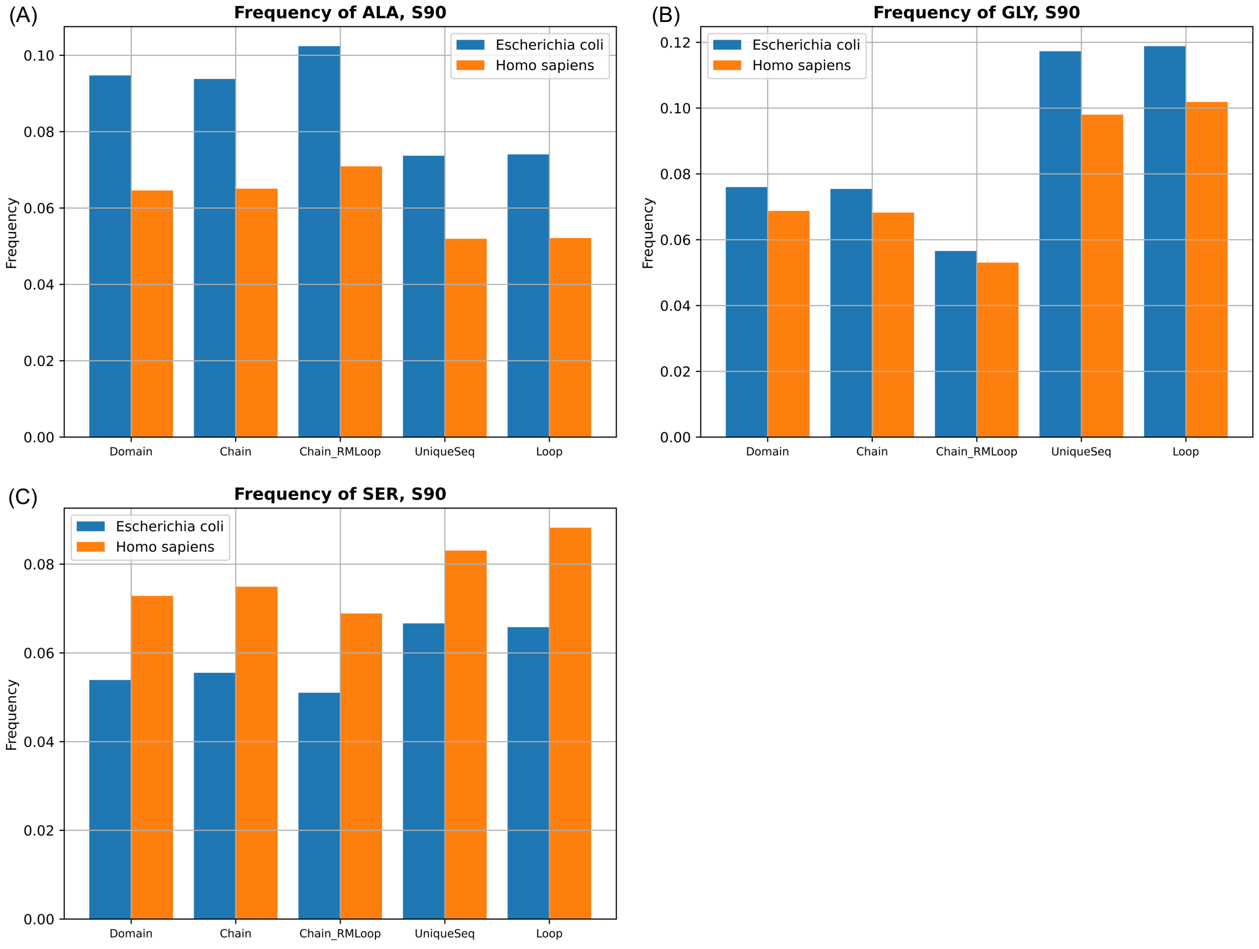


**Figure S6. Frequency of glycine, alanine, and serine in five datasets from humans and *E. coli.***

The datasets included only specific segments of amino acid sequences and were Domain (sequences of protein domain regions), Chain (sequences of whole protein chains), Chain_RMLoop (sequences of protein chains with loop region removed), UniqueSeq (all unique loop sequences), and Loop (all loop sequences, which indicated loop sequences are redundant).

(A)

| Redundancy | Protein chains | # of Loops | Annotated Loops | Unique LoopSeq | Non-redundant LoopSeq |
|---|---|---|---|---|---|
| All | 20,772 | 228,368 | 148,192 | 45,456 | 47.32% |
| S100 | 2,692 | 32,087 | 17,032 | 24,028 | 81.16% |
| S90 | 2,056 | 23,488 | 11,645 | 21,276 | 91.46% |

(B)

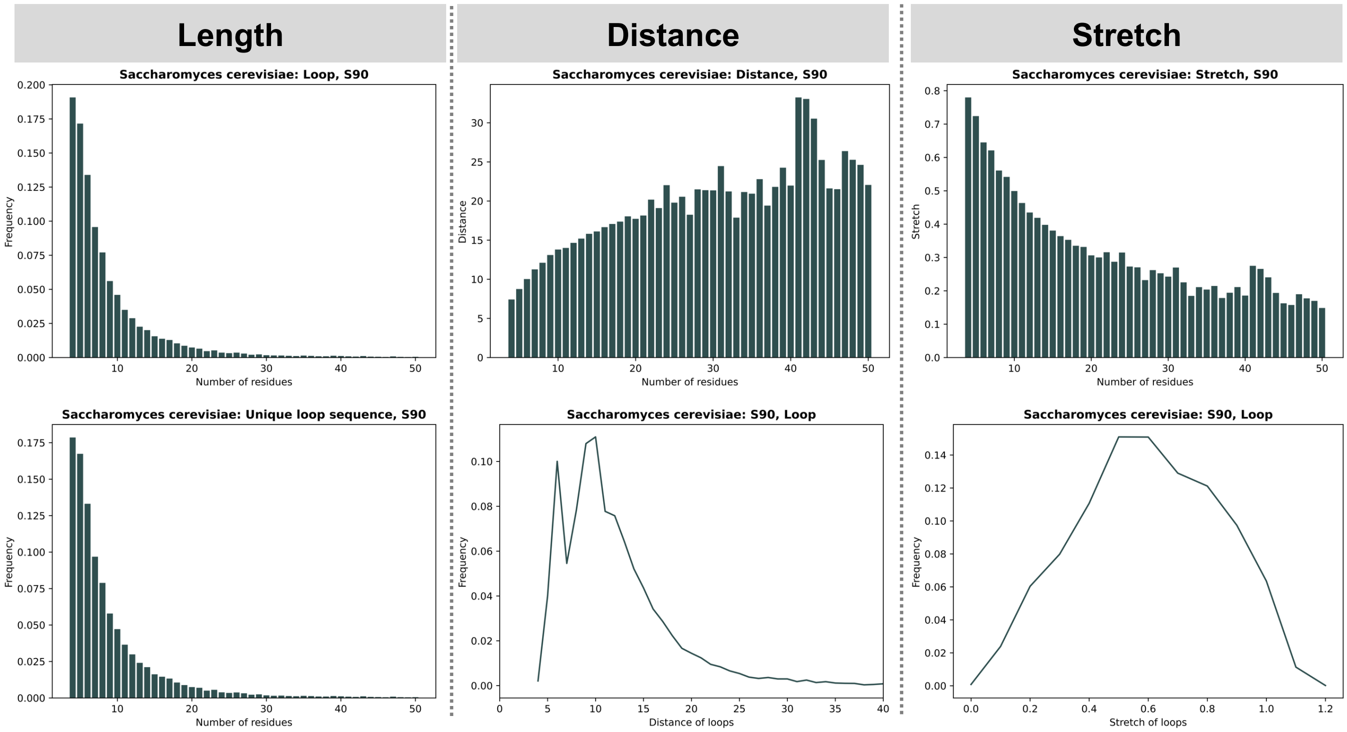


(C)

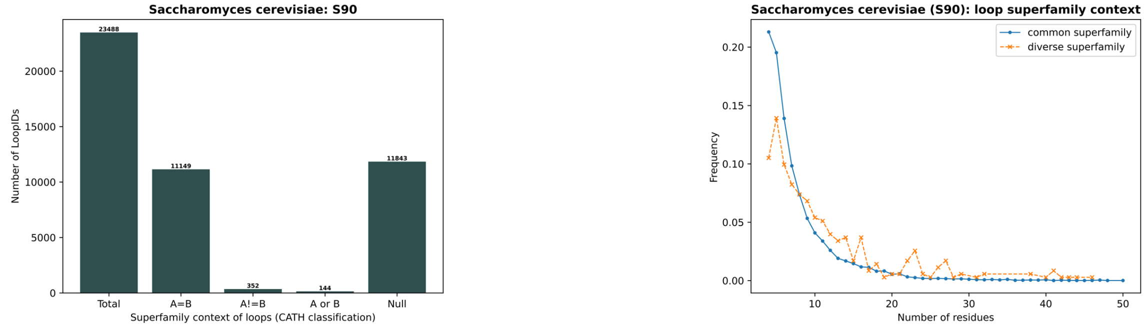


(D)

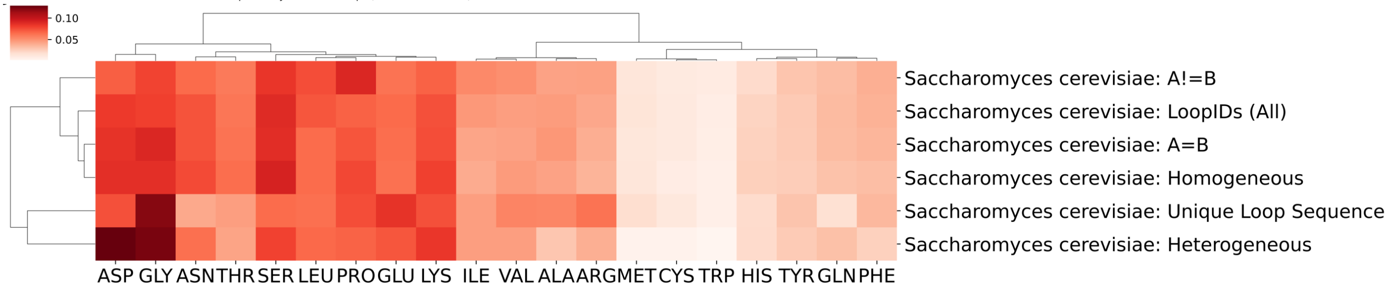


(E)

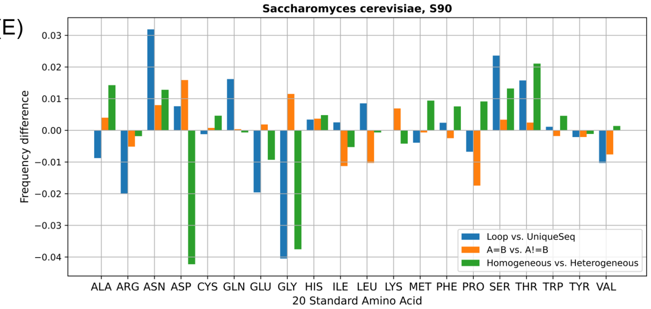


(F)

| # of redundant loop sequences | CATH_levels | Homogeneous | Heterogeneous |
|---|---|---|---|
| 789 | (C, A, T, H) | 703 | 86 |
| | (C, A, T) | 713 | 76 |
| | (C, A) | 719 | 70 |
| | (C) | 755 | 34 |

**Figure S7. Characterizing properties of *Saccharomyces cerevisiae* protein loops.**
(A) Protein loop statistics. (B) Visualizations of loop properties. (C) Superfamily context of a loop based on the CATH database . (D) Clustered heatmap of amino acid composition profiles. (E) Amino acid frequency differences among groups. (F) Statistics of superfamily context repertoire of loops from S90 non-redundant datasets.

(A)

| Redundancy | Protein chains | # of Loops | Annotated Loops | Unique LoopSeq | Non-redundant LoopSeq |
|---|---|---|---|---|---|
| All | 838 | 7,522 | 7,126 | 2,114 | 41.53% |
| S100 | 182 | 1,693 | 1,539 | 1,428 | 87.89% |
| S90 | 151 | 1,419 | 1,283 | 1,347 | 91.14% |

(B)

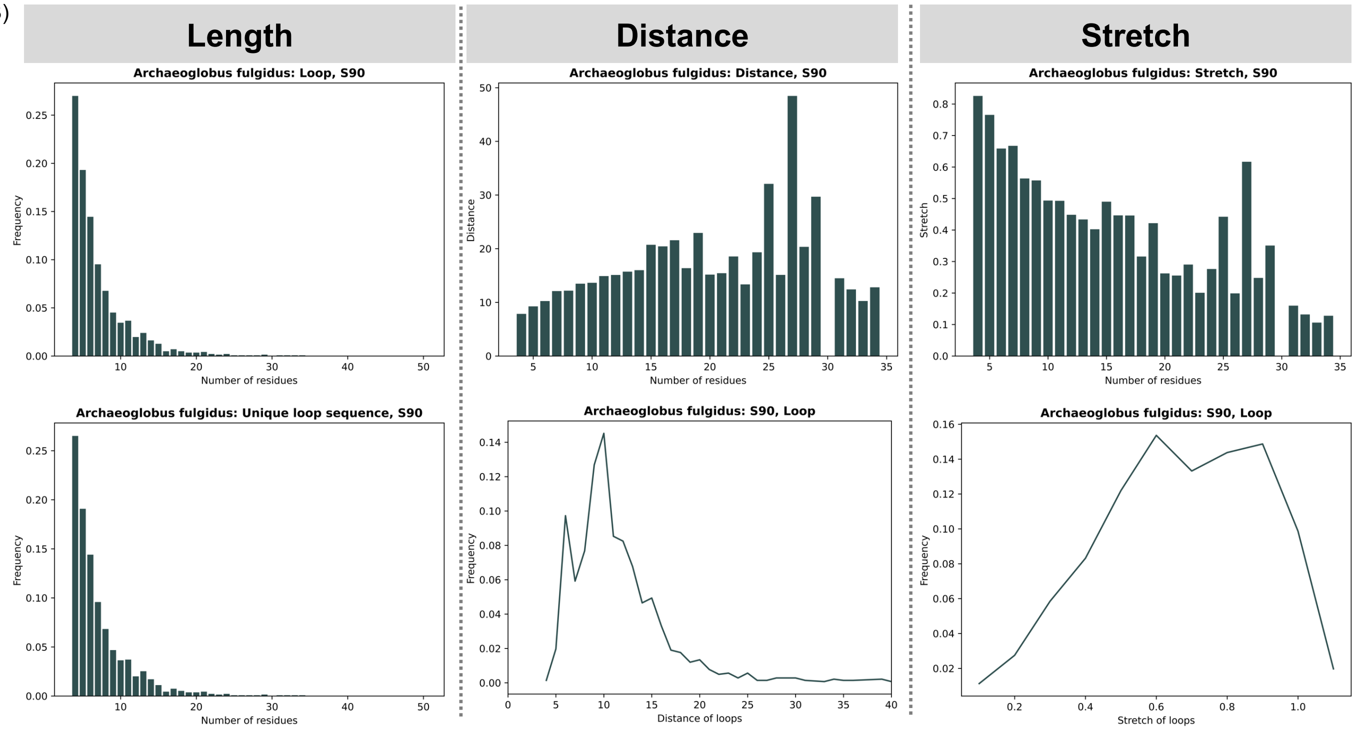


(C)

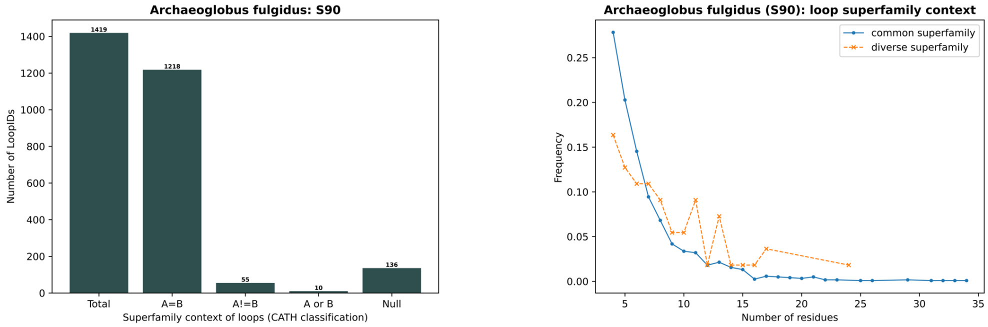


(D)

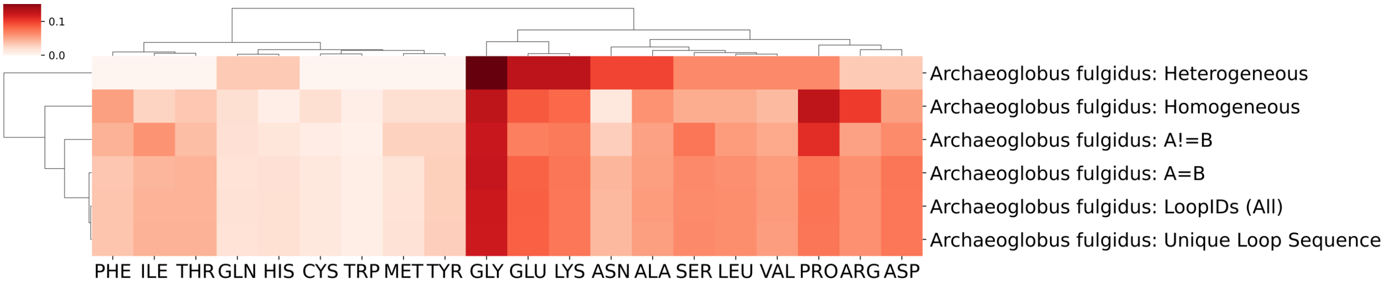


(E)

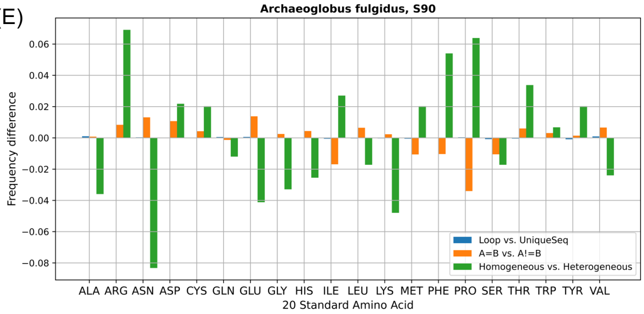


(F)

| # of redundant loop sequences | CATH_levels | Homogeneous | Heterogeneous |
|---|---|---|---|
| 789 | (C, A, T, H) | 24 | 5 |
| | (C, A, T) | 24 | 5 |
| | (C, A) | 25 | 4 |
| | (C) | 26 | 3 |

**Figure S8. Characterizing properties of *Archaeoglobus fulgidus* protein loops.**
(A) Protein loop statistics. (B) Visualizations of loop properties. (C) Superfamily context of a loop based on the CATH database. (D) Clustered heatmap of amino acid composition profiles. (E) Amino acid frequency differences among groups. (F) Statistics of superfamily context repertoire of loops from S90 non-redundant datasets.

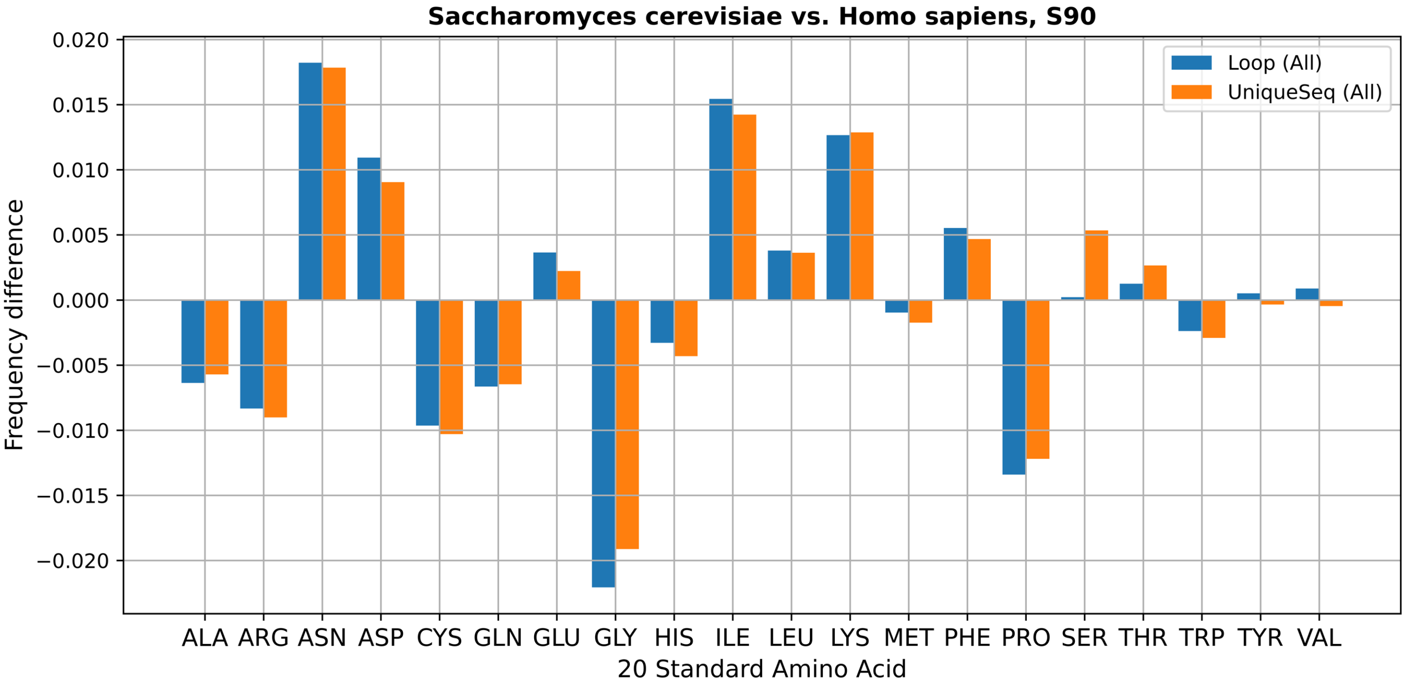


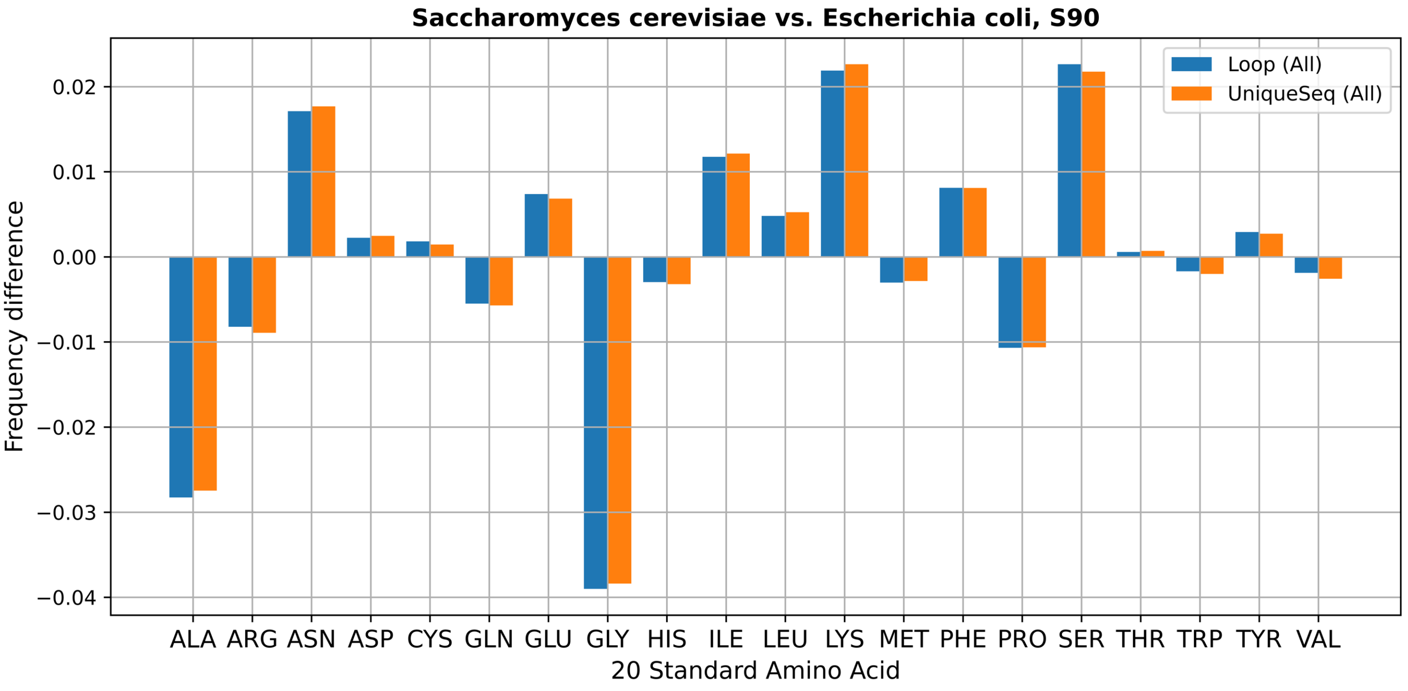


**Figure S9. Amino acid frequency comparisons between *Saccharomyces cerevisiae*, humans, and *E. coli*.** The human and E. coli frequencies were used as a baseline; that is, positive values indicate that the amino acids are more abundant in yeast.

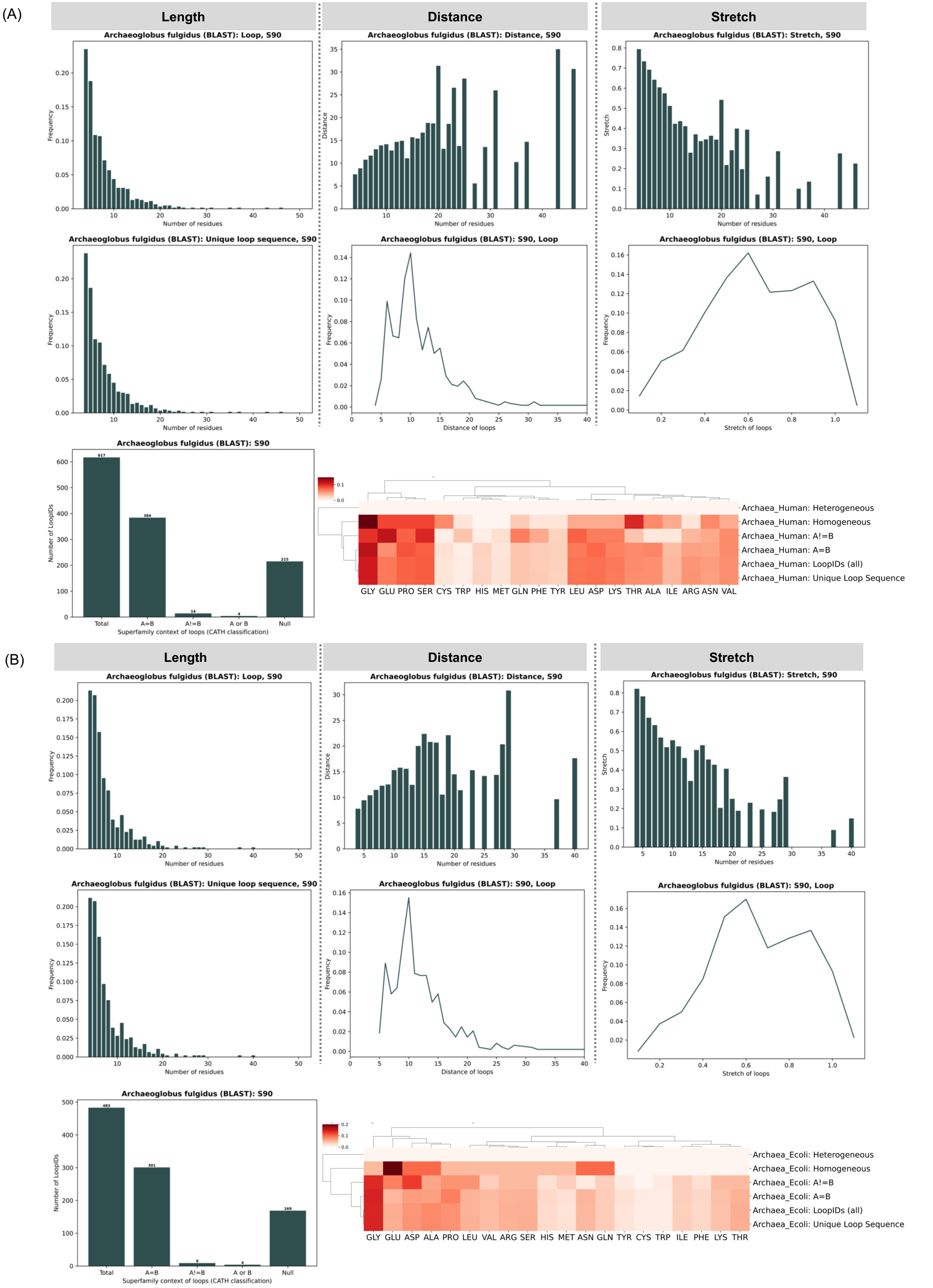


**Figure S10. Blast analysis using 151 protein chains from *Archaeoglobus fulgidus* against human and *E. coli* proteins.**

(A) and (B) represent the loop property characterizations using matched protein chains from human and E. coli, respectively. The following parameters were used to select the best matched amino acid sequence in the blast analysis: e value 1e-2 and the coverage of the query sequence (*Archaeoglobus fulgidus* chains) is ≥ 50%.

**Table S1. Top 30 conserved loop sequences from humans and *E. coli* and their comparisons to each other.**

**Ranking of Homo sapiens (S90)**

| ConservedSeq | Length | # of Loop (humans) | # of Loop (*E. coli*) |
|---|---|---|---|
| QPGGS | 5 | 102 | 1 |
| SSLR | 4 | 76 | 1 |
| DTEGY | 5 | 33 | 1 |
| YGLSENDEWTQDRAKPVT | 18 | 31 | 1 |
| DPQP | 4 | 30 | 1 |
| EQPALNDSR | 9 | 30 | 1 |
| YPDH | 4 | 28 | 1 |
| SSLT | 4 | 20 | 2 |
| TGGNSPV | 7 | 19 | 1 |
| GPPGTG | 6 | 19 | 2 |
| DFDSQTNVSQSKDSDV | 16 | 18 | 1 |
| GSKS | 4 | 17 | 1 |
| QNPRN | 5 | 17 | 1 |
| HPDK | 4 | 15 | 36 |
| RSMD | 4 | 15 | 1 |
| NKHMNADTD | 9 | 14 | 38 |
| GGGT | 4 | 13 | 2 |
| NKSDFA | 6 | 13 | 1 |
| SGLKPGV | 7 | 13 | 1 |
| NAASPN | 6 | 13 | 37 |
| YLLT | 4 | 13 | 37 |
| KPLG | 4 | 13 | 35 |
| MPNIPQ | 6 | 13 | 29 |
| AGAD | 4 | 12 | 10 |
| INGDKG | 6 | 12 | 37 |
| DKLY | 4 | 11 | 27 |
| VLPTFKGQPSKP | 12 | 11 | 20 |
| LGQNPT | 6 | 10 | 1 |
| LGTT | 4 | 10 | 2 |
| MFNLQE | 6 | 10 | 33 |

**Ranking of Escherichia coli (S90)**

| ConservedSeq | Length | # of Loop (humans) | # of Loop (*E. coli*) |
|---|---|---|---|
| NKHMNADTD | 9 | 14 | 38 |
| NAASPN | 6 | 13 | 37 |
| YLLT | 4 | 13 | 37 |
| INGDKG | 6 | 12 | 37 |
| HPDK | 4 | 15 | 36 |
| KPLG | 4 | 13 | 35 |
| MFNLQE | 6 | 10 | 33 |
| MPNIPQ | 6 | 13 | 29 |
| GRQT | 4 | 9 | 29 |
| DKLY | 4 | 11 | 27 |
| SGLLAEITPD | 10 | 8 | 24 |
| KDLLPNPPKT | 10 | 5 | 24 |
| VLPTFKGQPSKP | 12 | 11 | 20 |
| AKGK | 4 | 6 | 18 |
| ATGDGPD | 7 | 6 | 14 |
| TGDGPD | 6 | 3 | 13 |
| AGAD | 4 | 12 | 10 |
| KDLLPNPPKTWEE | 13 | 5 | 10 |
| SKVN | 4 | 2 | 10 |
| SAVN | 4 | 5 | 9 |
| SGLLAEITPA | 10 | 4 | 9 |
| SGRQT | 5 | 4 | 8 |
| AATGDGPD | 8 | 2 | 8 |
| ENGK | 4 | 3 | 7 |
| EGKT | 4 | 2 | 7 |
| LGVD | 4 | 1 | 7 |
| AADGG | 5 | 1 | 7 |
| TGGN | 4 | 3 | 6 |
| TPDGR | 5 | 2 | 6 |
| ADTP | 4 | 1 | 6 |

**Table S2. CATH classifications of the conserved loops in the A!=B group (S90).**

| Species | Chain | Length | Start position | Stop position | Sequence | CATH Start | CATH Stop |
|---|---|---|---|---|---|---|---|
| Homo_sapiens | 1DUG:A | 7 | 78 | 84 | HNMLGGC | 3.40.30.10 | 1.20.1050.10 |
| Homo_sapiens | 1I3K:A | 5 | 270 | 274 | GTGTG | 3.40.50.720 | 3.90.25.10 |
| Homo_sapiens | 1WSV:A | 4 | 53 | 56 | VSHM | 3.30.1360.120 | 3.30.70.1400 |
| Homo_sapiens | 2XSX:A | 4 | 126 | 129 | KGVP | 3.30.390.10 | 3.20.20.120 |
| Homo_sapiens | 3B97:A | 4 | 125 | 128 | KGVP | 3.30.390.10 | 3.20.20.120 |
| Homo_sapiens | 3E9L:A | 4 | 1919 | 1922 | LYDD | 3.30.420.230 | 1.20.80.40 |
| Homo_sapiens | 3FE1:A | 4 | 228 | 231 | THLG | 3.30.420.40 | 3.90.640.10 |
| Homo_sapiens | 3GDQ:A | 4 | 228 | 231 | THLG | 3.30.420.40 | 3.90.640.10 |
| Homo_sapiens | 3I33:A | 4 | 229 | 232 | THLG | 3.30.420.40 | 3.90.640.10 |
| Homo_sapiens | 3SOA:A | 5 | 301 | 305 | GAILT | 1.10.510.10 | 3.10.450.50 |
| Homo_sapiens | 3ZDX:B | 4 | 56 | 59 | FPVS | 3.30.1680.10 | 2.60.40.1510 |
| Homo_sapiens | 4EBC:A | 4 | 96 | 99 | GEDL | 3.40.1170.60 | 3.30.70.270 |
| Homo_sapiens | 4J8F:A | 4 | 226 | 229 | THLG | 3.30.420.40 | 3.90.640.10 |
| Homo_sapiens | 4JKH:A | 4 | 1919 | 1922 | LYDD | 3.30.420.230 | 1.20.80.40 |
| Homo_sapiens | 4PJ1:A | 4 | 373 | 376 | LSDG | 3.50.7.10 | 3.30.260.10 |
| Homo_sapiens | 4UM9:D | 4 | 60 | 63 | NPVS | 3.30.1680.10 | 2.60.40.1510 |
| Homo_sapiens | 4ZVX:A | 4 | 269 | 272 | LGGN | 3.40.605.10 | 3.40.309.10 |
| Homo_sapiens | 5DMZ:A | 4 | 782 | 785 | LGSK | 6.10.130.20 | 1.10.510.10 |
| Homo_sapiens | 5EVZ:A | 4 | 251 | 254 | THLG | 3.30.420.40 | 3.90.640.10 |
| Homo_sapiens | 5FPN:A | 4 | 228 | 231 | THLG | 3.30.420.40 | 3.90.640.10 |
| Homo_sapiens | 5PXJ:A | 4 | 759 | 762 | SGHQ | 3.30.40.10 | 1.20.920.10 |
| Homo_sapiens | 6ASY:A | 4 | 251 | 254 | THLG | 3.30.420.40 | 3.90.640.10 |
| Escherichia_coli | 1B8X:A | 7 | 78 | 84 | HNMLGGC | 3.40.30.10 | 1.20.1050.10 |
| Escherichia_coli | 1DKG:D | 4 | 225 | 228 | THLG | 3.30.420.40 | 3.90.640.10 |
| Escherichia_coli | 1DRU:A | 4 | 237 | 240 | ASSR | 3.30.360.10 | 3.40.50.720 |
| Escherichia_coli | 1Q16:B | 4 | 433 | 436 | VGLT | 1.10.3650.10 | 3.30.70.20 |
| Escherichia_coli | 1RKQ:A | 4 | 87 | 90 | TALS | 3.40.50.1000 | 3.30.1240.10 |
| Escherichia_coli | 1RLM:A | 4 | 82 | 85 | GELT | 3.40.50.1000 | 3.30.1240.10 |
| Escherichia_coli | 1TJ7:A | 6 | 103 | 108 | LHTGRS | 1.10.275.10 | 1.20.200.10 |
| Escherichia_coli | 1U6Z:A | 4 | 293 | 296 | SDGA | 3.30.420.150 | 3.30.420.40 |
| Escherichia_coli | 1U6Z:A | 4 | 310 | 313 | RHQD | 3.30.420.40 | 1.10.3210.10 |
| Escherichia_coli | 1XA3:A | 4 | 328 | 331 | LESN | 3.30.1540.10 | 3.40.50.10540 |
| Escherichia_coli | 1XVI:A | 4 | 184 | 187 | ASAG | 3.30.980.20 | 3.40.50.1000 |
| Escherichia_coli | 1YE9:A | 4 | 262 | 265 | IPRS | 1.10.10.1060 | 2.40.470.10 |
| Escherichia_coli | 1ZYL:A | 4 | 108 | 111 | SVGG | 3.30.200.70 | 1.10.510.10 |
| Escherichia_coli | 2RDZ:A | 4 | 444 | 447 | IGLN | 1.20.140.10 | 1.10.1130.10 |
| Escherichia_coli | 3A8J:A | 4 | 48 | 51 | VSHM | 3.30.1360.120 | 3.30.70.1400 |
| Escherichia_coli | 3ABS:B | 5 | 107 | 111 | LKEVP | 1.10.30.40 | 3.40.50.11240 |
| Escherichia_coli | 3JZ4:A | 4 | 255 | 258 | LGGN | 3.40.605.10 | 3.40.309.10 |
| Escherichia_coli | 3K5M:A | 4 | 37 | 40 | APQE | 2.40.50.590 | 3.30.70.2250 |
| Escherichia_coli | 3NZQ:A | 4 | 602 | 605 | EGDT | 2.40.37.10 | 1.10.287.3440 |
| Escherichia_coli | 4AZV:A | 4 | 310 | 313 | KLPG | 3.30.200.20 | 1.10.510.10 |
| Escherichia_coli | 4AZW:A | 4 | 310 | 313 | KLPG | 3.30.200.20 | 1.10.510.10 |
| Escherichia_coli | 4C4V:A | 4 | 421 | 424 | RNTG | 3.10.20.310 | 2.40.160.50 |
| Escherichia_coli | 4FZW:C | 4 | 202 | 205 | ATQP | 3.90.226.10 | 1.10.12.10 |
| Escherichia_coli | 4G9B:A | 4 | 94 | 97 | VLPG | 1.10.150.240 | 3.40.50.1000 |
| Escherichia_coli | 4JNE:A | 4 | 225 | 228 | THLG | 3.30.420.40 | 3.90.640.10 |
| Escherichia_coli | 6NPF:A | 4 | 125 | 128 | KGMP | 3.30.390.10 | 3.20.20.120 |

**Table S3. Descriptions of the top ten CATH superfamily and Pfam clan between humans and *E.coli*.**

| Species | Superfamily | Description | Pfam Clans | Description |
|---|---|---|---|---|
| Homo sapiens | 2.60.40.10 | Immunoglobulins | CL0011 | Immunoglobulin superfamily |
| Homo sapiens | 3.40.50.300 | P-loop containing nucleotide triphosphate hydrolases | CL0023 | P-loop_NTPase |
| Homo sapiens | 3.40.50.720 | NAD(P)-binding Rossmann-like Domain | CL0177 | Periplasmic binding proteins (PBPs) |
| Homo sapiens | 1.10.510.10 | Transferase(Phosphotransferase) domain 1 | CL0063 | NADP_Rossmann |
| Homo sapiens | 2.30.42.10 | PDZ domain | CL0016 | PKinase |
| Homo sapiens | 2.130.10.10 | YVTN repeat-like/Quinoprotein amine dehydrogenase | CL0186 | Beta_propeller |
| Homo sapiens | 3.40.190.10 | Periplasmic binding protein-like II | CL0159 | Ig-like fold superfamily (E-set) |
| Homo sapiens | 3.40.30.10 | Glutaredoxin | CL0466 | PDZ-like |
| Homo sapiens | 3.30.420.40 | ATPase, nucleotide binding domain | CL0029 | Cupin |
| Homo sapiens | 3.30.200.20 | Phosphorylase Kinase; domain 1 | CL0172 | Thioredoxin |
| Escherichia coli | 3.40.190.10 | Periplasmic binding protein-like II | CL0177 | Periplasmic binding proteins (PBPs) |
| Escherichia coli | 3.40.50.300 | P-loop containing nucleotide triphosphate hydrolases | CL0063 | NADP_Rossmann |
| Escherichia coli | 3.20.20.70 | Aldolase class I | CL0023 | P-loop_NTPase |
| Escherichia coli | 3.40.50.720 | NAD(P)-binding Rossmann-like Domain | CL0036 | TIM_barrel |
| Escherichia coli | 2.160.10.10 | Hexapeptide repeat proteins | CL0172 | Thioredoxin |
| Escherichia coli | 2.60.40.10 | Immunoglobulins | CL0536 | Hexapeptide repeat superfamily (HEXAPEP) |
| Escherichia coli | 3.40.30.10 | Glutaredoxin | CL0123 | Helix-turn-helix (HTH) |
| Escherichia coli | 3.40.50.150 | Vaccinia Virus protein VP39 | CL0061 | PLP_aminotran |
| Escherichia coli | 3.40.50.2300 | Response regulator | CL0021 | oligonucleotide/oligosaccharide binding (OB) |
| Escherichia coli | 3.30.565.10 | Histidine kinase-like ATPase, C-terminal domain | CL0186 | Beta_propeller |

### Table S4. Twenty standard amino acid compositions (ratios) of loops from various groups (S90).

| | Entire PDB | | | | | | Homo sapiens | | | | | | *Escherichia coli* | | | | | | Conserved Loops | | |
|---|---|---|---|---|---|---|---|---|---|---|---|---|---|---|---|---|---|---|---|---|---|
| | Loop (All) | Unique Seq. (All) | A=B | A!=B | Homogeneous | Heterogeneous | Loop (All) | Unique Seq. (All) | A=B | A!=B | Homogeneous | Heterogeneous | Loop (All) | Unique Seq. (All) | A=B | A!=B | Homogeneous | Heterogeneous | Conserved Loop Seq. | Related Loop (Human) | Related Loop (*E. coli*) |
| ALA | 6.29% | 6.25% | 6.37% | 6.40% | 5.76% | 6.68% | 5.21% | 5.19% | 5.21% | 5.33% | 4.88% | 5.10% | 7.40% | 7.37% | 7.40% | 8.04% | 7.14% | 9.91% | 6.78% | 6.12% | 7.28% |
| ARG | 4.63% | 4.72% | 4.58% | 4.76% | 4.57% | 4.42% | 4.99% | 5.09% | 4.93% | 4.72% | 5.03% | 4.66% | 4.98% | 5.08% | 4.94% | 5.22% | 5.12% | 4.06% | 4.26% | 3.94% | 3.33% |
| ASN | 5.95% | 5.88% | 5.90% | 5.03% | 6.19% | 6.93% | 5.49% | 5.50% | 5.55% | 4.44% | 6.00% | 6.01% | 5.60% | 5.52% | 5.63% | 4.70% | 5.79% | 5.32% | 6.06% | 5.69% | 6.73% |
| ASP | 7.72% | 7.63% | 7.87% | 6.53% | 7.65% | 9.61% | 7.08% | 7.18% | 7.13% | 6.62% | 7.46% | 8.41% | 7.95% | 7.84% | 8.08% | 7.09% | 8.74% | 8.78% | 9.54% | 9.38% | 9.66% |
| CYS | 1.34% | 1.39% | 1.29% | 1.27% | 1.68% | 0.60% | 2.10% | 2.17% | 1.95% | 2.75% | 2.14% | 1.29% | 0.96% | 1.00% | 1.05% | 0.87% | 0.93% | 0.20% | 0.51% | 0.30% | 0.33% |
| GLN | 3.46% | 3.53% | 3.38% | 3.33% | 3.36% | 2.96% | 4.08% | 4.08% | 4.06% | 3.90% | 3.89% | 3.31% | 3.96% | 4.00% | 3.94% | 4.09% | 3.91% | 3.13% | 3.33% | 4.09% | 3.26% |
| GLU | 5.89% | 6.00% | 5.82% | 6.24% | 5.34% | 6.25% | 6.17% | 6.34% | 6.04% | 6.75% | 5.92% | 6.39% | 5.80% | 5.87% | 5.72% | 5.98% | 5.79% | 5.59% | 5.71% | 5.13% | 4.84% |
| GLY | 11.13% | 10.46% | 11.69% | 10.05% | 11.78% | 15.21% | 10.19% | 9.80% | 10.64% | 9.69% | 10.86% | 16.62% | 11.88% | 11.73% | 12.29% | 9.46% | 12.58% | 20.08% | 18.49% | 18.06% | 16.93% |
| HIS | 2.45% | 2.55% | 2.50% | 2.36% | 2.71% | 1.82% | 2.64% | 2.75% | 2.64% | 2.42% | 2.64% | 1.46% | 2.61% | 2.64% | 2.77% | 2.53% | 2.71% | 1.60% | 1.75% | 1.42% | 1.80% |
| ILE | 3.56% | 3.72% | 3.45% | 3.92% | 3.34% | 2.85% | 3.18% | 3.36% | 3.08% | 3.29% | 3.02% | 2.04% | 3.55% | 3.57% | 3.50% | 3.81% | 3.46% | 4.32% | 2.26% | 1.71% | 2.46% |
| LEU | 6.47% | 6.52% | 6.25% | 7.24% | 5.93% | 5.97% | 6.81% | 6.88% | 6.60% | 7.18% | 6.35% | 6.72% | 6.71% | 6.72% | 6.38% | 7.90% | 5.44% | 5.85% | 7.02% | 7.32% | 7.83% |
| LYS | 5.83% | 5.77% | 5.73% | 5.76% | 5.57% | 6.55% | 6.16% | 6.11% | 6.18% | 5.87% | 6.16% | 6.92% | 5.24% | 5.13% | 5.12% | 4.42% | 5.25% | 5.59% | 5.81% | 5.40% | 6.98% |
| MET | 1.50% | 1.59% | 1.46% | 1.69% | 1.60% | 0.83% | 1.57% | 1.66% | 1.51% | 1.93% | 1.52% | 0.80% | 1.77% | 1.77% | 1.71% | 1.72% | 1.94% | 0.47% | 0.90% | 0.80% | 1.40% |
| PHE | 3.21% | 3.36% | 3.15% | 3.37% | 3.18% | 2.21% | 3.19% | 3.31% | 3.13% | 3.56% | 3.02% | 1.66% | 2.93% | 2.97% | 2.94% | 3.69% | 2.69% | 2.33% | 1.79% | 1.58% | 1.73% |
| PRO | 7.94% | 7.94% | 8.03% | 9.70% | 8.49% | 6.78% | 8.15% | 7.98% | 8.28% | 9.42% | 8.57% | 6.81% | 7.88% | 7.82% | 7.90% | 9.93% | 8.41% | 6.78% | 6.55% | 7.90% | 7.77% |
| SER | 7.85% | 7.62% | 7.77% | 7.03% | 7.73% | 7.90% | 8.82% | 8.31% | 8.98% | 7.68% | 8.45% | 10.18% | 6.58% | 6.66% | 6.48% | 6.70% | 5.92% | 6.18% | 8.06% | 9.88% | 6.63% |
| THR | 6.21% | 6.16% | 6.22% | 5.87% | 6.45% | 6.13% | 5.94% | 5.79% | 5.97% | 5.21% | 6.01% | 6.37% | 6.01% | 5.98% | 6.00% | 5.53% | 6.15% | 4.32% | 5.92% | 6.12% | 6.28% |
| TRP | 1.08% | 1.16% | 1.08% | 1.14% | 1.18% | 0.33% | 1.07% | 1.13% | 1.05% | 1.01% | 1.10% | 0.44% | 1.00% | 1.04% | 1.01% | 1.10% | 0.93% | 0.20% | 0.28% | 0.38% | 0.26% |
| TYR | 2.86% | 3.00% | 2.84% | 3.14% | 3.01% | 1.90% | 2.74% | 2.83% | 2.72% | 3.18% | 2.82% | 1.37% | 2.50% | 2.53% | 2.51% | 2.36% | 2.35% | 1.53% | 1.34% | 1.60% | 1.45% |
| VAL | 4.64% | 4.77% | 4.62% | 5.18% | 4.48% | 4.05% | 4.41% | 4.56% | 4.35% | 5.05% | 4.14% | 3.46% | 4.69% | 4.77% | 4.63% | 4.86% | 4.78% | 3.79% | 3.65% | 3.19% | 3.05% |

**Table S5. Loops with stretch values > 1.2. .**

| | Loop | | | Unique loop sequence | | |
|---|---|---|---|---|---|---|
| | Entire PDB | humans | *E. coli* | Entire PDB | humans | *E. coli* |
| 4_AA | 23 | 7 | 0 | 21 | 7 | 0 |
| 5_AA | 17 | 2 | 0 | 17 | 2 | 0 |
| 6_AA | 7 | 4 | 1 | 7 | 4 | 1 |
| 7_AA | 5 | 2 | 0 | 5 | 2 | 0 |
| 8_AA | 4 | 0 | 0 | 4 | 0 | 0 |
| 9_AA | 5 | 2 | 0 | 5 | 2 | 0 |
| 10_AA | 6 | 0 | 0 | 6 | 0 | 0 |
| 11_AA | 1 | 0 | 0 | 1 | 0 | 0 |
| 12_AA | 1 | 1 | 0 | 1 | 1 | 0 |
| 13_AA | 1 | 1 | 0 | 1 | 1 | 0 |
| 14_AA | 3 | 1 | 0 | 3 | 1 | 0 |
| 15_AA | 0 | 0 | 0 | 0 | 0 | 0 |
| 16_AA | 0 | 0 | 0 | 0 | 0 | 0 |
| 17_AA | 0 | 0 | 0 | 0 | 0 | 0 |
| 18_AA | 0 | 0 | 0 | 0 | 0 | 0 |
| 19_AA | 1 | 0 | 0 | 1 | 0 | 0 |
| 20_AA | 0 | 0 | 0 | 0 | 0 | 0 |
| 21_AA | 0 | 0 | 0 | 0 | 0 | 0 |
| Total | 74 | 20 | 1 | 72 | 20 | 1 |

**Table S6. Amino acid compositions of loops with larger stretches (> 1.2)**

| | Loop | | | Unique loop sequence | | |
|---|---|---|---|---|---|---|
| | Entire PDB | humans | *E. coli* | Entire PDB | humans | *E. coli* |
| ALA | 5.74% | 4.51% | 0.00% | 5.63% | 4.51% | 0.00% |
| ARG | 6.97% | 8.27% | 0.00% | 7.08% | 8.27% | 0.00% |
| ASN | 3.89% | 6.02% | 0.00% | 3.96% | 6.02% | 0.00% |
| ASP | 7.17% | 7.52% | 16.67% | 7.29% | 7.52% | 16.67% |
| CYS | 0.82% | 0.75% | 0.00% | 0.83% | 0.75% | 0.00% |
| GLN | 2.25% | 3.01% | 0.00% | 2.29% | 3.01% | 0.00% |
| GLU | 7.79% | 8.27% | 0.00% | 7.92% | 8.27% | 0.00% |
| GLY | 6.97% | 4.51% | 33.33% | 6.88% | 4.51% | 33.33% |
| HIS | 3.89% | 3.01% | 0.00% | 3.96% | 3.01% | 0.00% |
| ILE | 5.12% | 4.51% | 0.00% | 5.21% | 4.51% | 0.00% |
| LEU | 6.97% | 5.26% | 16.67% | 7.08% | 5.26% | 16.67% |
| LYS | 3.69% | 1.50% | 0.00% | 3.33% | 1.50% | 0.00% |
| MET | 2.66% | 3.01% | 16.67% | 2.50% | 3.01% | 16.67% |
| PHE | 1.84% | 2.26% | 16.67% | 1.88% | 2.26% | 16.67% |
| PRO | 9.22% | 8.27% | 0.00% | 9.17% | 8.27% | 0.00% |
| SER | 10.86% | 9.77% | 0.00% | 10.63% | 9.77% | 0.00% |
| THR | 7.17% | 6.77% | 0.00% | 7.29% | 6.77% | 0.00% |
| TRP | 0.41% | 0.75% | 0.00% | 0.42% | 0.75% | 0.00% |
| TYR | 1.64% | 2.26% | 0.00% | 1.67% | 2.26% | 0.00% |
| VAL | 4.92% | 9.77% | 0.00% | 5.00% | 9.77% | 0.00% |

**Table S7. Disorder statistics among the datasets.**

| Species | Redundancy | Total Loops | At least one disorder site | Fully disordered (except terminals) |
|---|---|---|---|---|
| Entire PDBs | all | 4,512,768 | 4.15% | 0.37% |
| | S100 | 905,728 | 4.44% | 0.40% |
| | S90 | 555,516 | 4.89% | 0.44% |
| Homo sapiens | all | 1,027,593 | 5.35% | 0.50% |
| | S100 | 192,496 | 5.64% | 0.49% |
| | S90 | 102,901 | 6.18% | 0.53% |
| Escherichia coli | all | 279,503 | 2.88% | 0.25% |
| | S100 | 49,331 | 3.18% | 0.26% |
| | S90 | 24,818 | 4.17% | 0.33% |
| Archaeoglobus fulgidus | all | 7,522 | 2.43% | 0.28% |
| | S100 | 1,693 | 3.07% | 0.41% |
| | S90 | 1,419 | 3.10% | 0.42% |
| Saccharomyces cerevisiae | all | 228,368 | 5.78% | 0.56% |
| | S100 | 32,087 | 8.78% | 1.27% |
| | S90 | 23,488 | 9.38% | 1.31% |
